# Non-blocking modulation as the major mechanism of sodium channel inhibition by riluzole

**DOI:** 10.1101/228569

**Authors:** AK Szabo, K Pesti, P Lukacs, MC Földi, Z Gerevich, B Sperlagh, A Mike

## Abstract

Modulated- or guarded receptor hypothesis; Channel block or modulation; “Voltage-sensor block” or “lipophilic block” are some of the questions that arise when drug effects on sodium channels are investigated. Understanding the exact mechanism of action for individual drugs is essential, because it is one of the major factors that determine their therapeutic applicability.

In this study we created a kinetic model of sodium channels, which can help us understand the modes of action for individual drugs in the context of these hypotheses. The model was constructed so that it could integrate the above hypotheses.

In particular we aimed to understand the mode of action of riluzole, a neuroprotective drug with a peculiar state-dependent inhibition. In experiments the inhibition by 100μM riluzole was full within the first two milliseconds, but it was almost completely removed between 2 and 20 ms post-depolarization. This abrupt loss of inhibition could not be explained by dissociation, which was proceeding with a time constant of ~300 ms. We propose that for sodium channel inhibitors binding without blocking is possible, and riluzole predominantly inhibits by non-blocking modulation. We used lidocaine as a reference compound, and found that non-blocking modulation, although less prominent, also may play a role in its mechanism of inhibition. Non-blocking modulation may selectively inhibit cells with pathological activity patterns, therefore this property may be a good trait to investigate in the development of sodium channel inhibitor drugs.

**SUMMARY:** Although never actually proven, presence of sodium channel inhibitors at their binding site is assumed to prevent ion conduction. Authors provide evidence from experiments and kinetic simulations that bound riluzole permits conduction and exerts its inhibitory effect almost entirely by modulation.

## INTRODUCTION

Local anesthetics, class I antiarrhythmics, and certain anticonvulsants are well-known examples of sodium channel inhibitors (SCIs; we avoid the term “blocker”, because that would convey an oversimplification of the inhibition mechanism, as we will see below). Compounds with SCI effect are currently being developed for several other indications (Eijkelkamp et al., 2012; England and de Groot, 2009; Tarnawa et al., 2007). Finding compounds with SCI property is in fact not a difficult task, voltage gated sodium channels are inhibited by a surprisingly large and diverse group of drugs (Huang et al., 2006; Lounkine et al., 2012; Zhang et al., 2015). However, potency alone does not ensure therapeutic effectiveness. The operation of sodium channels is essential for all excitable cells of the organism, therefore instead of indiscriminate inhibition, it would be desirable to selectively inhibit pathological activity. Fortunately most SCIs are inherently selective: they are more effective against specific conformations of the channel, mostly those that are formed upon depolarization. Several pathologies, such as inflammation, ischemic injury, traumatic injury, epilepsy, hyperalgesia, arrhythmias, *etc.,* can manifest themselves in hyperexcitability. Hyperexcitability is often caused by altered sodium channel function: an increased sensitivity to slightly depolarized membrane potential (~−65 to −50 mV), due to either left-shifted window current or enlarged persistent current component (Fischer et al., 2017; Morris et al., 2012). It is of great significance, therefore, that some of the SCI compounds are able to exert strong selectivity for channel conformations at this exact membrane potential range.

One striking example is the neuroprotective compound riluzole, which has a number of unique qualities: It was found very efficient in inhibiting the firing rate of neurons, with IC50 values ~0.5 to 4 μM (Centonze et al., 1998; Cho et al., 2015; Kuo et al., 2006; Urbani and Belluzzi, 2000). Its ability to inhibit the persistent component of the sodium current is within the same concentration range (Cho et al., 2015; Del Negro et al., 2002; Koizumi and Smith, 2008; Kuo et al., 2006; Ptak et al., 2005; Spadoni et al., 2002; Urbani and Belluzzi, 2000; Xie et al., 2011), which is not surprising, as the persistent sodium current is known to be crucial in action potential initiation. Although riluzole is considered to be a selective inhibitor of the persistent component(Cifra et al., 2013; Urbani and Belluzzi, 2000), it is important to note that it is equally potent in inhibiting transient currents evoked from slightly depolarized (~−65 to −50 mV) membrane potential range, as seen directly from the shifted steady-state availability curves within this range of membrane potentials (Benoit and Escande, 1991; Hebert et al., 1994; Lenkey et al., 2010; Ptak et al., 2005; Song et al., 1997; Urbani and Belluzzi, 2000), or from the calculated inactivated state affinities (Benoit and Escande, 1991; Hebert et al., 1994; Lenkey et al., 2010, 2011; Ptak et al., 2005), using the method of Bean et al. (Bean et al., 1983).

Ptak *et al.* (Ptak et al., 2005) argued that persistent component selectivity can be entirely explained by inactivated state preference. While the transient component is normally measured as the peak amplitude, i.e., within 1 ms after the voltage step, when the process of inactivation has not yet reached equilibrium; the persistent component is unavoidably measured in protocols which allow equilibration, either during a slow voltage ramp, or at the end of an extended voltage pulse. Indeed, when investigated under identical conditions no difference was found in potency for inhibiting transient and persistent components (Fig. 8E of Ptak et al., 2005).

At the same time, riluzole almost completely failed to inhibit resting sodium channels up to 10-30 μM: the inhibition was less than 10% when currents were evoked from hyperpolarized potentials (−160 to −120 mV) (Benoit and Escande, 1991; Hebert et al., 1994; Lenkey et al., 2010), and IC_50_ values calculated for resting inhibition ranged between 85 and 360 μM (Benoit and Escande, 1991; Cho et al., 2015; Hebert et al., 1994; Lenkey et al., 2010; Xie et al., 2011).

The inhibition also showed an extremely steep time-dependence: In a comparative study of 7 compounds (Desaphy et al., 2014), riluzole was the only one that showed no frequency-dependence between 0.1 and 10 Hz (at a holding potential of −120 mV), but a huge (~50-fold) increase in affinity at 50 Hz, −90 mV holding potential. This indicates that a 50-fold loss of affinity must occur within a time window of a few tens of milliseconds. Indeed, when the rate of recovery from inactivation was measured 10 μM riluzole produced almost full inhibition within the first 3-4 ms, but apparently lost its potency within ~30-40 ms (Benoit and Escande, 1991; Hebert et al., 1994). The fast offset kinetics must be matched with a fast onset kinetics as well, because the 20 ms of the first (conditioning) pulse in the recovery protocol was enough for reaching a complete inhibition.

What exact mechanism is behind this fast onset and recovery? Is it the association and dissociation of riluzole, or the conformational transitions of the channel? Is this fast kinetics a prerequisite for persistent component selectivity? Our aim in this study was to reach a more profound understanding of the mechanism of action riluzole exerts on sodium channels. The properties of riluzole clearly seem to set it apart among SCIs (Lenkey et al., 2010), and these special properties may be the reason of its remarkable therapeutic potency, most importantly as a neuroprotective agent, best studied in spinal cord injury (Fehlings et al., 2016; Nagoshi et al., 2015), but possibly also in other hyperexcitability-related disorders (Pittenger et al., 2008; Zarate and Manji, 2008). Understanding what exactly makes riluzole highly state-dependent and persistent selective may help developing more effective neuroprotective drugs.

Our approach in this study was that we first performed electrophysiology experiments: recorded the extent of inhibition under different conditions. The efficiency of SCI drugs profoundly depend on the temporal pattern of membrane potential, and individual drugs produce their own characteristic inhibition patterns. After recording these inhibition patterns for riluzole and lidocaine, therefore, we performed kinetic simulations, in which we tested which specific underlying mechanisms could account for the experimentally observed behavior.

We sought to use a model in which different theoretical concepts of sodium channel inhibitor mechanisms could be tested. Several different terminologies have been proposed for the explanation of experimentally recorded details of sodium channel inhibitor action, the best known concepts being the “modulated-” vs. “guarded receptor hypotheses”, “voltage-sensor block” vs. “lipophilic block”, and “channel block” vs. “modulation”. We aimed to construct a model that incorporates these concepts about SCI mechanisms, and thus one can characterize and test them in simulations. In the first section of this study we asked the question, how these hypothetical mechanisms of action would behave in standard experimental protocols. In the second section, we aimed to propose a plausible explanation for the peculiar behavior of riluzole, and also for lidocaine, which served as a reference compound.

We found that our experimental data could only be reproduced by supposing that the predominant mechanism of sodium channel inhibition by riluzole is almost pure modulation, without considerable channel block. We propose that riluzole inhibits sodium channels by stabilizing inactivated conformation, i.e., delaying recovery from inactivation. This means that riluzole binding in itself does not prevent conduction: once channels have recovered to riluzole-bound-resting state they can be reactivated, and riluzole-bound-open channels can conduct ions.

## METHODS

### Cell line generation and cell culture

For studies on WT Na_V_1.4 channels, stable transfection of Chinese hamster ovary (CHO) cells was performed. The coding sequence of the rat Na_V_1.4 sodium channel was inserted into a modified pBluescript KS (Stratagene) vector (pCaggs IgG-Fc) capable to recombine to the murine Rosa26 BAC (Zboray et al., 2015). Briefly, Na_V_1.4 was inserted into the vector at *AscI* sites under the control of the Caggs promoter. Recombination to the *Rosa26* BAC was carried out by Recombineering (Muyrers et al., 1999; Zhang et al., 1998). Original BAC clone RP24-85I15 was derived from the BACPAC Resources Center (Children’s Hospital Oakland Research Institute). Na_V_1.4 BAC was transfected into CHO DUKX B11 (ATCC CRL-9096) suspension cells by Fugene HD (Promega) transfection reagent according to the manufacturer’s recommendations. Cell clones with stable vector DNA integration were selected by the addition of G418 antibiotic to the culture media (400 mg/ml) for 14 days.

CHO cells were maintained in Iscove’s Modified Dulbecco’s Medium with 25mM HEPES and L-Glutamine (Lonza) supplemented with 10% v/v fetal calf serum, 200mM L-glutamine, 100 U/ml of penicillin/streptomycin, 0.5 mg/mL Geneticin(Life Technologies, Carlsbad, CA, USA) and 2% ProHT Supplement (Lonza). Cells were plated onto 35 mm Petri dishes and cultured for 24-36 hours.

For studies on binding site mutant channels, transient transfection of tsA201 cells was performed as described previously (Lukacs et al., 2014). Cells were grown in Dulbecco’s modified Eagle’s medium supplemented with 10 % fetal bovine serum and 100 U/ml penicillin/streptomycin (Life Technologies) A mixture of plasmids coding 1.5 μg of F1579A mutant r Na_V_1.4 α subunit, 0.2 μg of sodium channel β1 subunit, and 0.02 μg of eGFP were transiently transfected into tsA201 cells in 35-mm dishes using TurboFect Transfection Reagent (Thermo Fisher Scientific Baltics UAB, Vilnius, Lithuania) according to the manufacturer’s instructions. Sixteen hours later, cells were dissociated from the dish surface by treatment with a 0.25% trypsin solution (Life Technologies) for approximately 2 minutes, pelleted, resuspended in growth medium, and were kept for 24 to 72 hours before recording.

### Electrophysiology

Borosilicate patch pipettes with resistances between 1.5 and 3 MΩ were pulled and filled with intracellular solution consisting of (in mM): 55 CsCl, 65 CsF, 10 NaCl, 10 EGTA, 10 HEPES, adjusted to pH 7.3 with CsOH (~5 mM). The extracellular solution consisted of (in mM): 140 NaCl, 5 KCl, 2 CaCl_2_, 1 MgCl_2_, 5 HEPES, adjusted to pH 7.3 with NaOH. Osmolality of intra- and extracellular solutions was set to ~290 and ~300 mOsm, respectively. Only cells with series resistance < 9 MΩ were used. Data were sampled at 20 kHz, and filtered at 5 kHz. Currents were recorded using an Axopatch 200B amplifier and the pClamp software (Molecular Devices). We used three different voltage protocols: The voltage-dependence of availability was studied using a standard “steady-state inactivation” (**SSI**) protocol; the time course of recovery of availability (“repriming”) was studied using a standard “recovery from inactivation” (**RFI**) protocol, and finally, the possible difference between repriming and inhibitor dissociation was studied using a protocol that consisted of repeated three-pulse trains (**3PT**), and a period of drug application. The protocols are described in detail and illustrated in the Results section (experimental sub-section). For drug application we used the “liquid filament switch” method using 1.5 mm outer diameter theta glass tubes (Harvard Apparatus, Holliston, MA) and a Burleigh LSS-3200 ultrafast solution switching system (for details see Pesti et al., 2014). Solution reservoirs were connected to the pressure control unit of a DAD-12 solution exchange system (ALA Scientific Instruments Inc., Farmingdale, NY), this allowed optimization of flow rate. Solution exchange was complete within 1-4 ms. Riluzole and lidocaine were obtained from Sigma.

### Modeling

Simulations are based on numerically solving a set of differential equations each describing the occupancy of a conformational state:

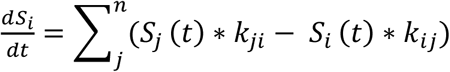

where *S_i_(t)* is the occupancy of a specific state at time *t*, and *S_j_(t)* is the occupancy of a neighboring state, *n* is the number of neighboring states; *k_ij_* and *k_ji_* are the rate constants of transitions between neighboring states. Differential equations were solved during simulations using a fourth-order Runge-Kutta method. We used the Berkeley Madonna v8.3.18 software (www.berkeleymadonna.com/).

We constructed a Markov equivalent of a Hodgkin-Huxley-type model, with two voltage-dependent gates corresponding with activation/deactivation and fast-inactivation/recovery-from-fast-inactivation. Slow inactivated states were not included in this model for the sake of simplicity. The two gates operated independently, so their position defined four drug-free states: **R**esting (**R**), **O**pen (**O**), **C**losed-Fast-Inactivated (**C**) and Open-**F**ast-Inactivated (**F**) (Fig.1A). Voltage-dependence of rate constants was defined by three-parameter exponential equations:

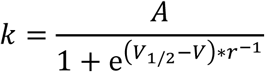

where the three free parameters *A, V_1/2_* and *r* defined the limiting value, the inflection point and the maximal slope, respectively. These parameter values (Table S1) have been previously optimized (Karoly et al., 2010) and we only needed a minor adjustment to match the kinetics and voltage-dependence of r Na_V_1.4 channels.

**Figure 1.**
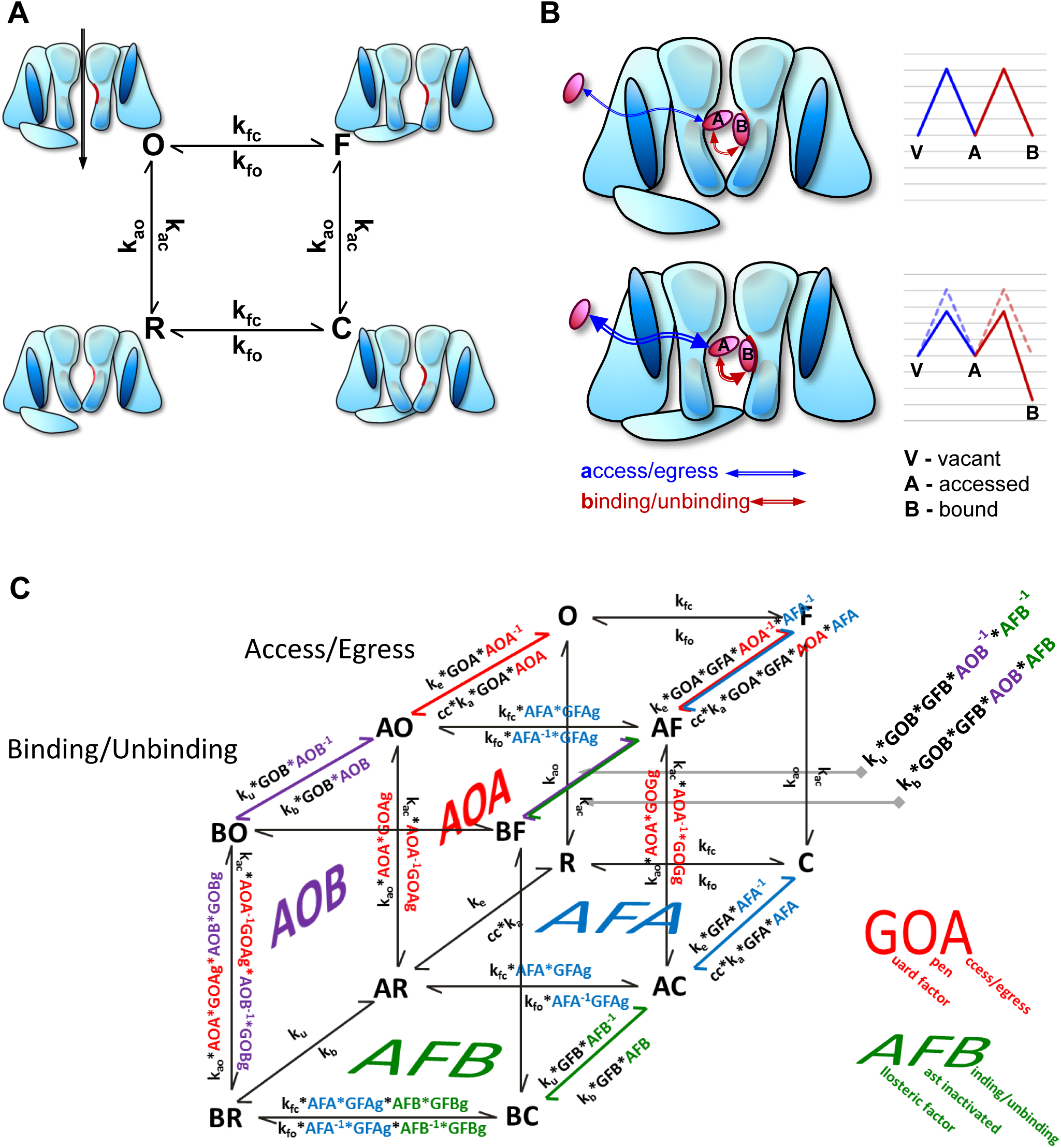
Structure of the model. (A) Schematic representation of the Hodgkin-Huxley type Markovian model without drug-accessed or drug-bound states. (B) Schematic drawings and energy diagrams illustrating the sub-processes of association and dissociation. Upper panel illustrates **R** (resting) conformation where access, egress, binding and unbinding all proceed with the same rate, 1000 s^-1^. Gridlines on the energy diagram indicate 1 kcal/mol. Lower panel illustrates **F** (inactivated) conformation, where accessibility is 10-fold increased (GFA = 10), and affinity is 100-fold increased by both 10-fold accelerated binding and 10-fold slowed unbinding (**AFB** = 10, **GFB** = 1). On the energy diagram dashed lines in pastel colors show resting state energy diagram for comparison. (C) Structure of the model, and calculation of rate constants using allosteric and guard factors. Parameter values are given in Table S2.

We aimed to study the concepts “modulated-” and “guarded receptor hypothesis”, as well as “voltage-sensor block” and “lipophilic block” by kinetic modeling. For this reason it seemed best to divide the processes of association and dissociation into two sub-processes: access/egress indicates entry/exit of the drug molecule into the inner vestibule of the channel, while binding/unbinding describes forming and breaking of chemical bonds between the ligand and its binding site. Figure 1B upper panel illustrates the case when energy barriers of all four processes are equal, therefore they all proceed with the same rate, which in this case was chosen to be 1 ms^-1^. (One might argue that the binding reaction must be much faster, but this describes binding to resting conformation, and in fact we found that reproducing experimental data required that we supposed a substantial energy barrier here.) Changing the conformation of the channel is expected to affect both sub-processes, as illustrated schematically in Fig. 1B, lower panel: in this case upon inactivation both accessibility and affinity are increased. These processes were modeled by adding drug-accessed (**AX**; meaning that the drug has entered the inner vestibule) and drug-bound (**BX**) states to all four states, where “**X**” may be any of the drug-free states: **R**, **O**, **C** or **F** (Fig.1C). Instead of association and dissociation rate constants, we defined access and egress rate constants (**k_a_**, and **k_e_**), as well as binding and unbinding rate constants (**k_b_**, and **k_u_**). Similarly, instead of a dissociation constant, we defined the “egress constant” (**K_E_** = **k_e_**/**k_a_**) and the “unbinding constant” (**K_U_** = **k_u_**/**k_b_**). For **AX** states, state-dependent accessibility was introduced by defining “accessibility factors” or “guard factors” (**GOA** and **GFA** for open and fast inactivated states, respectively), which altered both access and egress rates depending on the position of the activation or the inactivation gate. (The first character of factors indicates whether it is an **A**ffinity or a **G**uard factor, the second whether it refers to the **F**ast inactivated or **O**pen state, and the third one whether it regards to the **A**ccess/egress or the **B**inding/unbinding step.) For example energy diagrams in Fig. 1B (blue lines) illustrate the case when **GFA** = 10, which means that both access to- and egress from inactivated states are tenfold faster. For **BX** states state-dependent affinity was introduced by defining “affinity factors” (**AFB** and **AOB**) which altered either or both of binding and unbinding rates. Energy diagrams in Fig. 1B (red lines) illustrate the case when **AFB** = 10, furthermore, both rates are equally affected, which means that binding to inactivated state is tenfold faster, unbinding from inactivated states is tenfold slower, therefore, affinity to inactivated states is hundredfold higher.

Although in the simplest case the state dependence of the first step (access/egress) is modified by the guard factors only, while in the second step (binding/unbinding) by the affinity factors only; there is a possibility in the model to allow guard factors and affinity factors for both steps. This allows us to define if the conformational transition (e.g. inactivation) should affect binding, unbinding or both (Fig. 1C). For example when **GFB** = 1, rates of binding and unbinding are equally affected, like in the energy diagram shown in Fig. 1B. When **GFB** = **AFB**, only the binding rate, while when **GFB =** 1**/AFB**, only the unbinding rate is affected. In simulations **GFB** = 1/**AFB** was assumed as a default.

In general, affinity factors determine the relative free energy level of drug-accessed and drug-bound states, as compared to the free energy level of vacant states. Guard factors determine the overall rate of specific transitions (access/egress, binding/unbinding, as well as gating transitions), and at the same time they determine the balance between onward and backward rates: whether difference in free energy level is due to altered onward rates, altered backward rates, or both. While affinity factors must be the same for binding reactions and gating reactions to maintain detailed balance, guard factors obviously can be different (for example it is possible that in inactivated state affinity is increased by decreased unbinding rate (k_u_), with unchanged binding rate (k_b_); while both gating transitions: inactivation and recovery from inactivation are changed). For this reason we had to introduce separate “guard factors of gating” (Fig. 1C). For example when **GFBg** = 1, the rate of inactivation and recovery from inactivation are equally affected. When **GFBg = AFB**, only the rate of inactivation, while when **GFBg** = 1/**AFB** (which was the default value in simulations), only the rate of recovery is affected.

### Data analysis

Experimental as well as simulated traces were imported to Microsoft Excel, where curve fitting was performed using the Solver add-in. Steady-state availability (**SSI**) curves were fit by the following form of the Boltzmann function: *I(V)/I_max_* = (1+exp(*V-V_1/2_*)/*k*)^-1^, where *I_max_* is the maximal current, *V_1/2_* is the voltage of half availability, and *k* is the slope factor.

Recovery (**RFI**) curves were fit with mono- or bi-exponential functions: *I(t_ip_)/I_max_* = *A_1_*\*[1-exp(-*t_ip_*/*τ_1_*)]*^x^* + *A_2_*\*[1-exp(-*t_ip_*/*τ_2_*)], where *I_max_* is the maximal current (evoked by the first pulse), *t_ip_* is the length of the interpulse interval, *A_1_* and *A_2_* are the fractions of the amplitude that the two components contribute, *τ_1_* and *τ_2_* are time constants of the two components, and *x* is an exponent, which we included in the function, because we found that allowing it to differ from unity greatly improved fits. Its value ranged between 1.3 and 3.3, the best fit was obtained with x = 2.24 ± 0.17 for control curves. The time constant of the best fit function is dependent on the exponent: for example when exponential fitting on the first power gave 1 ms as a time constant, fitting the same data with an exponential on the second power gave 0.59 ms, and on the third power 0.45 ms. For the sake of comparability, therefore we fixed the value of the exponent at *x* = 2 in most cases.

Onset and offset of drug effect in the **3PT** protocol were fit with single exponential functions. The time constants obtained this way could be used to calculate the extent of inhibition expected, if a single-step binding process was supposed. In a single-step binding process time constants of onset and offset can be calculated from association and dissociation rate constants the following way: *τ_on_* = 1/(*k_a_*+*k_d_*), *τ_off_* = 1/*k_d_*, and inhibition can be calculated as: *Inh* = *k_a_*/(*k_a_*+*k_d_*), therefore the uninhibited fraction can be calculated as: (1-*Inh*) =*τ_on_* /*τ_off_*. As we will discuss below, riluzole did not bind in a single-step process, therefore this calculation did not correctly give the uninhibited, but rather the unbound fraction.

The concept of apparent affinity has been expounded previously (Lenkey et al., 2011), it is a useful way to represent the potency of SCIs under specific circumstances. The potency of SCIs can change radically even on the sub-millisecond time scale, depending on the temporal pattern of membrane potential, due to the dynamics of state-dependent accessibility and affinity. By calculating apparent affinity values these changes in potency can be monitored and compared. Apparent affinities (*K_app_*) can be calculated from the simplified Hill equation: when one-to-one binding is assumed, the Hill equation is reduced to *Inh* = *cc*/(*cc* + *K_app_*), where *Inh* is the inhibited fraction and *cc* is the drug concentration. From this *K_app_* is calculated (for example suppose that 100 μM of a compound causes the inhibited fraction to be 0.2, 0.5 or 0.8, the apparent affinity would then be 400 μM, 100 μM and 25 μM, respectively). The calculation is most accurate at ~50 % inhibition, but becomes increasingly inaccurate as inhibition approaches either 0 or 100 %.

In figures 1, 4, 5, and 7 we show energy diagrams of access/egress and binding/unbinding. The height of energy barriers was calculated from the Arrhenius equation: *k = A*exp(-E_a_/RT*), where k is the rate constant, *E_a_* is the activation energy, and *A* is the frequency factor, which *e.g.* in the case of access could mean the frequency of trials of the molecule to enter the inner vestibule. Since we do not have information regarding this, we arbitrarily chose a value that was faster than all rates in our simulations, (10^6^ s^-1^), and expressed energy barriers as relative to this value. Gridlines indicate 1 kcal/mol relative to this arbitrary value. Tenfold change in a rate constant corresponds with 1.36 kcal/mol energy difference.

Data are presented as arithmetic mean ± SEM, including the range of values whenever it is relevant, and the number of cells tested (n). For pairwise comparisons, paired, two-tailed Student’s *t* test was used, preceded by logarithmic transformation when necessary. In each case the exact value of p is given.

### Online supplemental material

Table S1, available as a Microsoft Excel file, shows parameter values for all models discussed in the study. The worksheet also helps to visualize energy diagrams for each model, and for any other combination of parameters.

## RESULTS

### Theoretical background - The modulated- and guarded receptor hypotheses

In order to properly understand the concepts that underlie our simulation studies, it is necessary to briefly summarize the two major hypotheses, and recent experimental data supporting their predictions.

#### *The modulated receptor hypothesis* (MRH)

(Hille, 1977; Hondeghem and Katzung, 1977) has been the first major hypothesis for the explanation of the characteristic properties of SCI action; such as frequency-, use-, and membrane-potential-dependence. According to this model, inactivation alters the conformation of the drug binding site, resulting in a higher affinity. If the affinity is changed by gating (*i.e.,* inactivation), it follows that gating rates (inactivation and recovery from inactivation) also must change upon drug binding. However, this change in gating rates cannot be monitored directly, by measuring conductance, because modulation usually coexists with channel block, and since drug-bound channels are blocked anyway, their modulation remains hidden. By observing indicators of molecular conformation other than the conducted current (e.g. monitoring the movement of voltage sensors by gating current measurement or by using fluorescent probes) it has been proven that energetics of gating is indeed altered by drug binding (Arcisio-Miranda et al., 2010; Hanck et al., 1994; Liu et al., 2002; Muroi and Chanda, 2009; Sheets et al., 2011).

#### *The guarded receptor hypothesis* (GRH)

(Starmer et al., 1984), has been proposed as the major alternative of MRH. This theory argues that the characteristic features of inhibition (voltage-, and frequency-dependence, as well as shift of the availability curve) can be explained without the assumption of altered gating. According to the GRH, it is not the binding site’s affinity that is altered by conformational change, but the site’s accessibility to the drug. In the most widely used variant of the model, “case 2”, the drug can only access the binding site when the activation gate is open, while the position of the inactivation gate also potently influences accessibility (fast access/egress with open inactivation gate, and slow access/egress when the gate is closed). Because access and egress are equally affected by the position of the inactivation gate, detailed balance is maintained in the model.

#### The conceptual difference between MRH and GRH is not the question of inactivated vs. open state, but of affinity vs. accessibility to specific states

Although the MRH focuses on the inactivated state, while the GRH discusses open state as an example, the essential difference is not which state is more important in drug effects, but whether the essence of the mechanism is conformation-dependent affinity or conformation-dependent accessibility. The principles of MRH (increased affinity) can be applied to open conformation, and the principles of GRH (increased accessibility) to inactivated conformation without difficulty, as we will show below.

#### The GRH has an MRH element hidden within

Although the guarded receptor hypothesis argues that characteristic properties of inhibition (frequency-dependence, holding potential dependence, shift of the availability curve, etc.) can be explained without assuming that bound drug molecules modify gating, it in fact supposes constrained gating of drug-bound channels (e.g. in “case 2” the activation gate of drug-bound channels cannot close), which can be conceived as an extreme case of the modulated receptor mechanism: If we began with the MRH, and supposed much higher affinity to channels with their activation gate open (so that practically there would be no binding to channels with closed activation gates), furthermore supposed strongly modified gating (so that bound drug molecules practically would not allow activation gates to close), then what we would get is exactly “case 2” of the GRH (Starmer et al., 1984).

#### Structural correlates and experimental proofs

Since the proposal of these two alternative hypotheses, much has been learned about the possible physical correlates of these concepts, especially within the last few years. Several bacterial sodium channel structures have been solved, representing different conformational states of the channel; these may resemble different conformations of eukaryotic sodium channels (Clairfeuille et al., 2016). These structures have been used in molecular dynamics studies in order to learn about possible access pathways and binding sites. It is becoming increasingly clear that the MRH vs. GRH problem is not an either-or question: Both mechanisms must contribute to state-selectivity of drug molecules, because both altered access pathways and altered binding sites have been observed in these conformations. Most recently two eukaryotic sodium channel structures have been solved (Shen et al., 2017; Yan et al., 2017). The differences between the closed NavPaS structure and the open-inactivated EeNaV1.4 structure confirm fundamental changes in both accessibility and affinity. The physical correlate of accessibility is either the activation gate (for permanently charged compounds), or for most SCI compounds it is most probably one of the fenestrations, which also change their structure with conformational transitions of the channel (Bagneris et al., 2014; Catterall and Swanson, 2015; Lukacs et al., 2014; Yan et al., 2017). The state-dependent changes in affinity are best proven by the fact that mutations of key residues involved in use-dependent (i.e., mostly inactivated state) inhibition, do not prevent tonic (i.e., mostly resting state) inhibition (Ragsdale et al., 1994). In addition, it has been shown that the key residue of the binding site, a phenylalanine in domain IV, S6 (“DIVS6 Phe”; F1764 in Nav1.2) does not only affect affinity, but also relays drug effect to the voltage sensor domains, thereby affecting gating (Hanck et al., 2009; Muroi and Chanda, 2009).

### Theoretical background – “rapid block” vs. “discrete block” and “lipophilic block” vs. “voltage-sensor block”

The question of “*rapid block*” vs. “*discrete block;*” as well as “*lipophilic block*” vs. “*voltage-sensor block*” are apparently unrelated to the MRH vs. GRH problem. However, as we will see, the three terminologies might have some conceptual overlap, and therefore in kinetic modeling the same topology can be used for investigating these concepts.

Two forms of block have been observed when the effects of lidocaine (Zamponi and French, 1993) or the quaternary amine lidocaine analog QX-314 (Gingrich et al., 1993) were studied on single channel sodium currents: rapid block (manifested as a reduction of unitary current amplitudes) and discrete block (shown by distinct several-millisecond-long closing events). Mutation of the key DIVS6 Phe abolished discrete block, but did not alter rapid block (Kimbrough and Gingrich, 2000). The same mutation has also been shown to abolish “voltage-sensor block”, i.e., the effect of SCIs on voltage sensors as observed in gating current measurements, and the shift of steady-state availability curves (Hanck et al., 2009). This suggests that “discrete block” might not be only block in the strict sense (i.e., inhibiting conduction sterically or electrostatically), but may as well involve inhibition by modulation (i.e., by stabilization of a non-conducting conformation). The voltage-independent and mutation-insensitive component of the inhibition was termed “lipophilic block”.

We see some common elements in all three terminologies, and therefore built our model to be able to accommodate all three: ligand-channel interaction occurs in two steps, the first step being the entry into the inner vestibule, and the second step the actual binding. The rate of the first step is much determined by the conformation of the channel, whether the entry occurs through the fenestrations (hydrophobic pathway (Hille, 1977)), or through the activation gate (hydrophilic pathway (Hille, 1977)). Increased accessibility is supposed to equally accelerate both entry to- and exit from the inner vestibule, thus fulfilling the requirements of the GRH. In addition, because discrete block required the DIVS6 Phe, we can suppose that the presence of the drug within the inner vestibule without binding to the binding site in itself only causes rapid block. Finally, because the effect on voltage sensors also required the DIVS6 Phe; either when gating charge immobilization was studied (Hanck et al., 2009), or when movement of the voltage sensor was monitored by voltage-clamp fluorimetry (Muroi and Chanda, 2009), we can also conclude that the inhibitor being within the inner vestibule but not bound to the binding site cannot produce “voltage-sensor block”, only “lipophilic block”. (The term may not be the most appropriate, considering that the permanently charged, and therefore not lipophilic QX-314 was also able to cause a certain extent of block in the mutant channels (Muroi and Chanda, 2009)).

The second step of actual binding involves an interaction of the ligand with the DIVS6 Phe (and possibly other residues). We suppose that this must be a higher energy interaction, because it is responsible for most of the affinity of SCIs. The presence of discrete block is consistent with this, because a higher energy interaction should take more time to disconnect. A high energy binding should also be required to change the energetics of conformational equilibria of the whole channel as supposed by the MRH, thus causing the characteristic phenomena abundantly documented in the literature of SCIs: leftward shift of the steady-state availability curve, delayed recovery from inactivation, use-dependence, frequency-dependence, etc.; as well as the more explicitly described alterations in the movement of voltage sensors revealed by recording gating charges(Hanck et al., 1994, 2009), or fluorescent signals(Arcisio-Miranda et al., 2010; Muroi and Chanda, 2009).

### Experiments – Steady-state availability

In an earlier comparative study of SCIs we have noticed that riluzole showed a peculiar mode of action with an extremely high state-dependence (Lenkey et al., 2010). The major properties of inhibition by riluzole are shown in Fig. 2.

**Figure 2.**
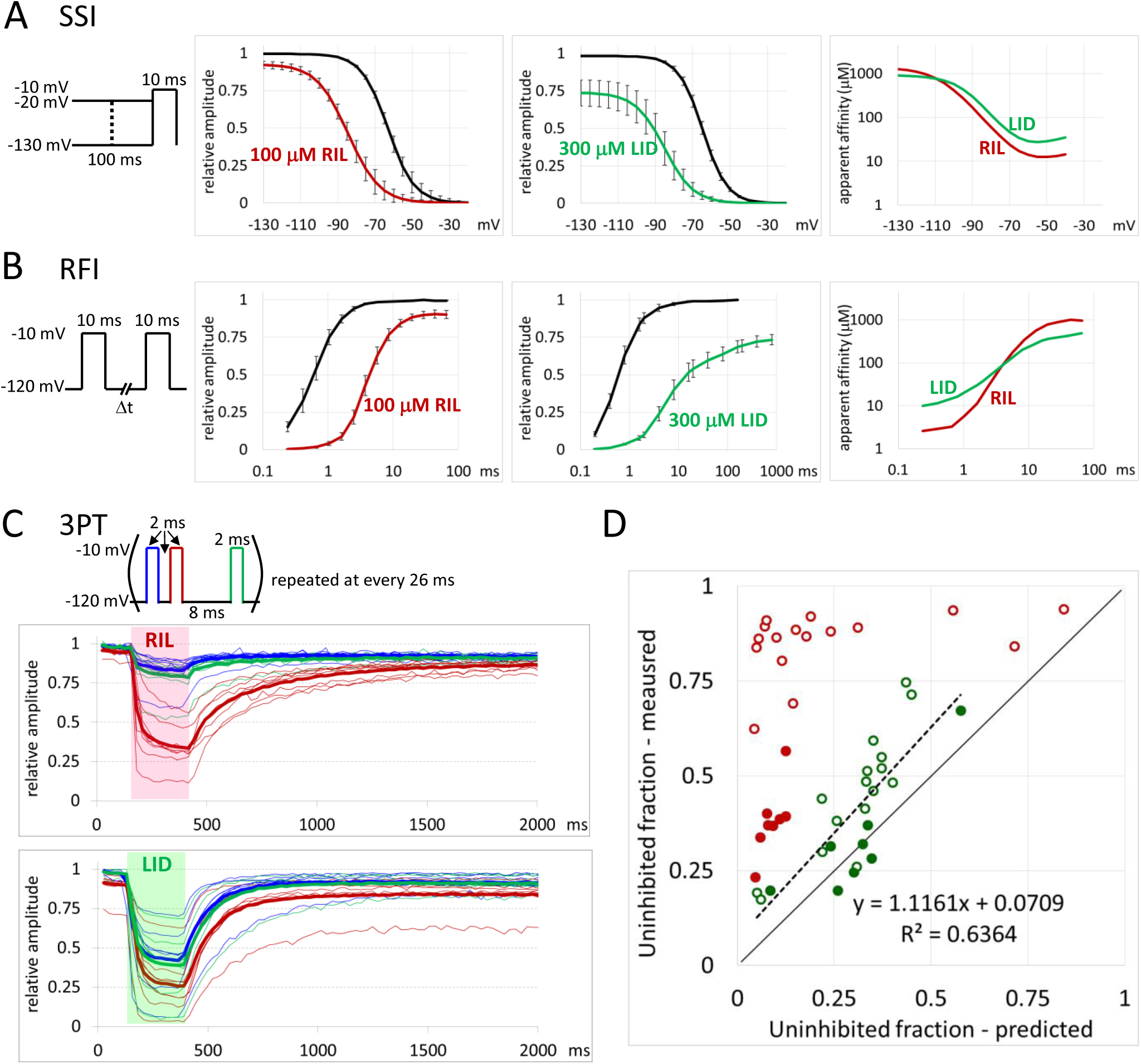
The effect of riluzole and lidocaine on WT channels. (A) The SSI protocol investigates availability of channels as a function of pre-pulse potential. Peak amplitudes of sodium currents were normalized to the maximal control amplitude of the same cell. Apparent affinities were calculated for each pair of amplitudes (drug/control), as described in Methods. (B) The **RFI** protocol investigates availability of channels as a function of interpulse interval. Peak amplitudes of sodium currents were normalized to the maximal control amplitude of the same cell. Apparent affinities were calculated for each pair of amplitudes (drug/control), as described in Methods. (C) The **3PT** protocol was designed to compare recovery of availability in the presence of the drug (within-train recovery during drug perfusion), and recovery of availability upon removal of the drug (between-train recovery after drug application). Blue, red, and green lines indicate peak amplitudes of currents evoked by the 1^st^, 2^nd^, and 3^rd^ pulse, respectively. Thin lines show data from individual cells (n = 8 for both drugs), thick lines show averages. Shaded areas indicate the 260 ms long drug pulse. (D) Comparison of calculated unbound fraction and measured uninhibited fraction. For each evoked current uninhibited fraction was measured after subtracting the slow inactivation component. Onset and offset time constants were determined by mono-exponential fitting. Unbound fraction was calculated from time constants as described in Methods. Red and green circles indicate the effect of riluzole and lidocaine, respectively. Open circles show time constants from 1^st^ and 3^rd^ pulse-evoked current amplitudes, filled circles show data from 2^nd^ pulse-evoked currents. For lidocaine the equation of the regression line (dashed line) and the R^2^ value are also shown. Thin black line indicates the case when calculated unbound fraction equals measured uninhibited fraction.

In the steady-state inactivation (**SSI**) protocol, its effect (Fig. 2A) is minimal resting state inhibition (0.92 ± 0.024 of control; range: 0.84 to 1.01; n = 7), but a strong leftward shift of the steady-state availability curves (V_1/2_ values were shifted by −23.5 ± 1.3 mV; range: −18.7 to −26.9 mV). In order to evaluate the extent of effect at different holding potentials, we converted inhibition values into apparent affinity values (see Methods). The apparent affinity value increased hundredfold, from 1261 μM at −120 mV holding potential to 12.5 μM at −40 mV. In the case of lidocaine, the extent of shift was similar (-24.2 ± 3.96, n = 6), but resting inhibition was stronger (0.74 ± 0.086). For this reason, the state-dependence was somewhat lower, its apparent affinity values changed from 899 μM to 27.5 μM within the −120 to −40 mV membrane potential range.

### Experiments – Recovery from inactivation

The difference in apparent affinity was even more striking in the case of the “recovery from inactivation” (**RFI**) protocol (Fig. 2B). At <1 ms interpulse intervals the inhibition by 100 μM Riluzole was almost full; the apparent affinity was 2.56 to 5.87 μM, however, it rapidly decreased until it reached ~1000 μM at >40 ms interpulse intervals. There was a hundredfold decrease between 1 and 10 ms. Recovery curves both in control and in the presence of riluzole could be better fit with a single exponential equation on the 2nd to 3rd power (the exponent was 2.24 ± 0.17 for control, and 2.81 ± 0.23 for riluzole n = 6). As we discuss in Methods, the exponent was fixed at 2.0 for the sake of comparability, in which case the time constant for control was 0.53 ± 0.09 ms, and it was increased by riluzole 6.51-fold (range: 5.53- to 10.34-fold) to 3.44 ± 0.39 ms. Recovery in the presence of 300 μM lidocaine, on the other hand, could only be fitted with a bi-exponential equation, where best fits gave an exponent of 1.54 ± 0.13 (n = 6) for the fast component, while the slow component was on the first power. For the sake of comparability we used a bi-exponential equation where the fast component was on the 2nd power, in this case the time constant was increased from 0.52 ± 0.06 ms (control) 7.83-fold (range: 6.76- to 9.46-fold), to 4.09 ± 0.53 ms in the presence of lidocaine (n = 6). The fast component contributed 69.5 ± 4.7 % of the amplitude (range: 52.3 to 87.7%), the slow component was 103.5 ± 12.4 ms (range: 73.9 to 155 ms), and it was responsible for the remaining 30.5 ± 4.7 % of the amplitude. As in the case of riluzole, apparent affinities for lidocaine decreased with increasing interpulse interval from 10.3 μM to 535 μM, although this 52-fold decrease was much less than the 391-fold decrease observed in the presence of riluzole.

Delayed repriming (recovery of availability during hyperpolarization) is conventionally interpreted as gradual dissociation of the drug, or as stabilization of a slow inactivated conformation (Gawali et al., 2015; Karoly et al., 2010). Recovery and dissociation are interrelated: bound inhibitors delay recovery (by stabilizing inactivated state), and delayed recovery hinders dissociation (because inactivated state has higher affinity). Drug-bound resting state is generally considered blocked, therefore not available for activation, furthermore, if resting affinity is low, this state should also be short-lived, and rapidly terminated by dissociation. To interpret the results with riluzole we needed to ask if drug-accessed and drug-bound resting states (**RA** and **RB** in the model; see Fig. 1C) might be activatable, *i.e.,* drug-accessed and drug-bound open states (**OA** and **OB**) might conduct.

### Experiments – The three-pulse-train protocol

To investigate the sub-processes of dissociation, and to help interpreting the experimentally observed recovery during the **RFI** protocol, we designed a simple three-pulse-train protocol (**3PT**) (see Fig. 2C). The control pulse was followed by a second pulse after a 2 ms hyperpolarizing gap, and then by a third pulse after an 8 ms gap. The 2 ms gap was chosen because in the absence of riluzole this time is enough for most channels to recover, but in the presence of riluzole most channels are still inhibited. The 8 ms gap is only four times longer, but already allows recovery of the majority of channels even in the presence of 100 μM riluzole. We studied the onset and offset of drug effect using a theta-tube fast solution exchange system, by which the solution around the cell can be changed within a few milliseconds. This was fast enough for this protocol, because the total length of the three-pulse train was 16 ms, after which the cells were kept for 10 ms at −120 mV resting potential before the next train. This high frequency (trains were evoked at every 26 ms) caused a minor decrease in amplitude along the experiment due to slow inactivation, but this compromise was necessary to have sufficient temporal resolution.

The results of short perfusion of either 100 μM riluzole or 300 μM lidocaine are shown in Fig. 2C. After six control trains, at 160 ms after the start of the protocol, drug perfusion was started, which was finished at 400 ms (drugs were perfused during 9 trains). Recovery was observed for 62 trains (1612 ms). Thin lines show the plot of peak amplitudes for 8 – 8 individual cells both in the case of riluzole and lidocaine. First, 2^nd^ and 3^rd^ pulse-evoked amplitudes are shown in blue, red and green, respectively. Thick lines show averages. Shaded areas indicate the interval of drug perfusion. Onset and offset time constants were determined by single exponential fitting, after subtraction of the slow inactivation component. For the peaks evoked by the 2^nd^ pulses of each train, the onset time constant for riluzole was 27.7 ± 5.37 ms (range: 9.95 to 49.2 ms), while the offset time constant was 310 ± 52.7 ms (range: 185 to 540 ms) (n = 8). Onset time constants for the 1^st^ and 3^rd^ pulses were similar (within the same range), but offset time constants were somewhat faster (for 1^st^ pulse-evoked currents the time constant was 137 ± 29.7 ms; range 52 to 303 ms; which was 2.44 ± 0.46 times faster than for 2^nd^ pulse-evoked currents; p = 0.0048). For lidocaine both onset and offset time constants were similar for all three pulse-evoked currents, for the 2^nd^ pulses the values were 40.7 ± 3.92 ms (range: 20.4 to 52.1 ms), and 148 ± 17.73 ms (range: 81.4 to 240 ms), respectively (n = 8).

In the case of riluzole it was obvious that the large difference between the 2^nd^ and 3^rd^ pulse-evoked amplitudes, which reflects the recovery seen in the **RFI** protocol, could not be due to full dissociation (unbinding – egress – partitioning to aqueous phase), because, even after the drug had been washed out the inhibition was still present in the 2^nd^ pulses of the next, as well as several following trains. Within-train recovery proceeded with a time constant of 6.11 ± 0.70 ms (when fitted with an exponential on the first power, see Methods), while between-trains recovery proceeded with a time constant of 310 ± 52.7 ms. The fast time constant may be interpreted as representing modulated gating, while the slow time constant as representing unbinding. In the case of lidocaine, the difference between the 1^st^ and 2^nd^ pulse-evoked amplitudes was much smaller, which is consistent with the RFI curve being less steep within this range. In addition, the time constant of the between-trains recovery (138.5 ± 11.2 ms) was similar to the slow time constant of the **RFI** curves (103.5 ± 12.4 ms, see above), therefore the same process might be reflected in both. This is unlike in the case of riluzole, where **RFI** curves showed no signs of a detectable slow component. For the **3PT** traces, we calculated the extent of binding from onset and offset time constants (see Methods), and the calculated unbound fraction radically differed from measured uninhibited fraction (Fig. 2D), suggesting that observable inhibition severely underestimates the actual occupancy of binding sites. In contrast, in the case of lidocaine the unbound fraction calculated from onset and offset time constants relatively well predicted the uninhibited fraction, that was directly measured (see Fig. 2D) The linear correlation between calculated unbound fraction and measured uninhibited fraction was highly significant (R=0.80, p=3.03*10^-6^). In summary, the discrepancy between observable extent of inhibition and the time constants suggests the presence of a hidden, non-blocking but riluzole-bound fraction of channels. The occupancy of binding sites can only be detected when the modulation of gating is monitored. We supposed that the fast within-train recovery represented modulated gating and the slow between-train recovery represented unbinding. However, as in the case of riluzole, this would imply that drug-bound channels are also available for activation, and can produce about half the conductance of vacant channels. We, therefore, need to consider alternative explanations as well: the fast process might have represented modulated gating and unbinding, while the slow process drug egress; it might even be possible that the fast component represented modulated gating, unbinding and egress, while the slow component membrane-to-aqueous-phase partitioning only. We tried to decide this question by experiments with binding site mutant channels.

### Experiments – The F1579A binding site mutant

The DIVS6 phenytoin residue (F1579 in Na_V_1.4 channels) has been shown to be crucial in high affinity binding to depolarized conformations. We reasoned that mutation of this residue must affect binding/unbinding processes, while it should have minor effect on access/egress, and none on partitioning. We therefore performed experiment on the F1579A mutant channels. Result of experiments are shown in Fig. 3. For both riluzole and lidocaine the strong inhibition seen in the **3PT** protocol was radically reduced, although not fully abolished, and, importantly, the slow between-trains recovery disappeared (Fig. 3C). Onset and offset time constants became almost equally fast, approaching the time resolution of the protocol, which prevented the accurate calculation of inhibition from time constants (Fig. 3D; time constants were between 20 and 60 ms for both drugs, for both onset and offset, and for all three pulses). The mutation did not eliminate the apparent inconsistency between unbound fraction, calculated from time constants, and the uninhibited fraction, measured directly, in fact it was now evident in the case of lidocaine as well. This might indicate the existence of secondary binding sites. The shift of the **RFI** curves was also much reduced (Fig. 3C) for both drugs. The fast component became dominant, it contributed 91.2 ± 3.9 % and 91.5 ± 3.1 % of the amplitude in the case of riluzole and lidocaine, respectively. The prolongation of the fast component was minimal, only 1.24- fold in the case of riluzole (from 0.466 ± 0.041 to 0.553 ± 0.045 ms; n = 6), and 1.12-fold in the case of lidocaine (from 0.404 ± 0.029 to 0.456 ± 0.045 ms; n = 7). (As with wild type data, the fast component of the bi-exponential equation was on the second power for comparability.) The slow component was also much faster than in wild type: 4.44 ± 1.62 ms for riluzole, and 4.08 ± 0.66 ms for lidocaine.

**Figure 3.**
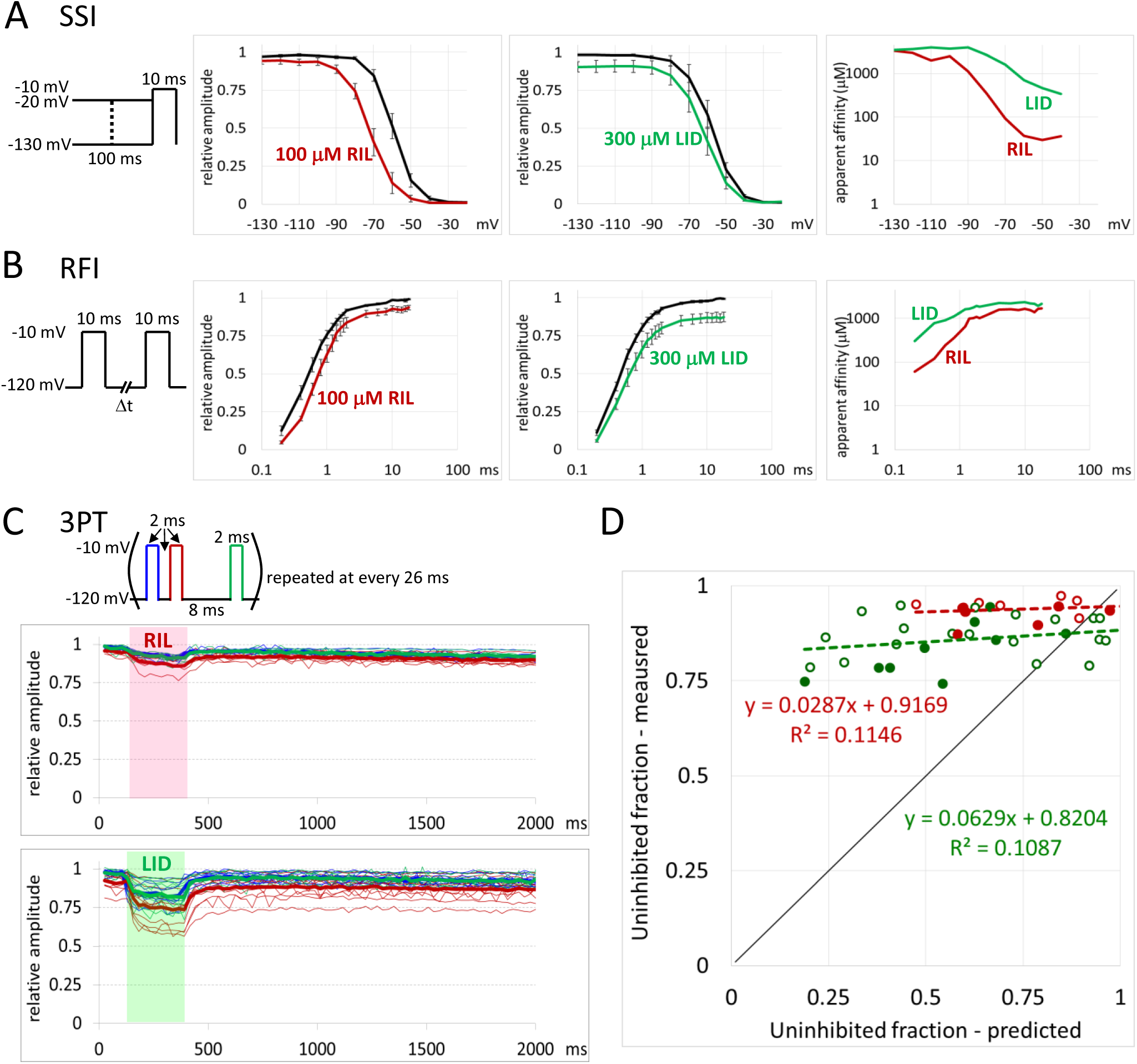
The effect of riluzole and lidocaine on F1579A mutant channels. (A) **SSI** protocol. and (B) **RFI** protocol. Peak amplitudes of sodium currents were normalized to the maximal control amplitude of the same cell. Apparent affinities were calculated for each pair of amplitudes (drug/control), as described in Methods. (C) **3PT** protocol. Blue, red, and green lines indicate peak amplitudes of currents evoked by the 1^st^, 2^nd^, and 3^rd^ pulse, respectively. Thin lines show data from individual cells (n = 6 for riluzole, n = 7 for lidocaine), thick lines show averages. Shaded areas indicate the 260 ms long drug pulse. (D) Comparison of calculated unbound fraction and measured uninhibited fraction. The slow inactivation component was subtracted before determining onset and offset time constants by mono-exponential fitting. Unbound fraction was calculated from time constants as described in Methods. Red and green circles indicate the effect of riluzole and lidocaine, respectively. Open circles show time constants from 1^st^ and 3^rd^ pulse-evoked current amplitudes, filled circles show data from 2^nd^ pulse-evoked currents. Equations of regression lines and the R^2^ values are also shown.

In spite of the much reduced inhibition in the **3PT** and **RFI** protocols, riluzole was still able to shift **SSI** curves considerably in mutant channels (-12.1 ± 2.71 mV), and this was the only protocol where it showed a relatively high apparent affinity at depolarized membrane potentials (29 to 36 μM at membrane potentials −60 to −40 mV) (Fig. 3A). A more thorough understanding of the fine details of inhibition by riluzole and lidocaine in mutant channels will require further experiments. The main conclusions of our experiments with mutant channels are the following: The F1579 residue was required for modulation as seen in the **RFI** protocol, and also for the slow between-trains recovery as seen in the **3PT** protocol. The most likely explanation of these is that this residue is crucial in high affinity binding, and therefore unbinding is a slow process, which is reflected in the slow between-trains recovery. At the same time, bound drugs allow a certain degree of conduction, which is hindered in the case of lidocaine, but almost unhindered in the case of riluzole.

### Simulations – Aims of kinetic modeling

The shift of the availability curve and the delayed recovery without considerable resting state inhibition, as well as the inconsistency between time constants and extent of inhibition were easy to detect, but proved difficult to explain. The significance of solving this puzzle is that SCI compounds with these specific properties are likely to have wide therapeutic index, *i.e.,* significant efficacy against pathological hyperexcitability without much inhibition of normal physiological function. Based on these properties of inhibition we hypothesized that riluzole must be able to bind to sodium channels and modify their gating without blocking them. In order to test if this mechanism was possible, and to understand this hypothetical mechanism in quantitative details, we intended to build a kinetic model which we could use to test different hypotheses regarding mechanisms of action.

Our aim of course was not only to understand the peculiar mechanism of riluzole, but also to construct a general kinetic model of sodium channels that is able to accommodate major hypotheses for the modes of action of SCIs. We intended to test and compare hypothetical modes of action. What type of inhibition would be caused by the MRH, the GRH, or a mixture of the two? What type of inhibition would be caused by block, modulation, or the mixture of the two? What mechanism could be the basis of “voltage-sensor block” and “lipophilic block”? Can we explain why mutation of the DIVS6 Phe decreases the shift in the steady-state availability curve?

We also intended to create a kinetic model by which different mechanisms of various compounds could be simulated, and plausible mechanisms could be proposed for them. We intended to address questions such as which conformations are preferred, how much this preference is due to increased accessibility (as proposed by the GRH) and to increased affinity (as proposed by the MRH), and what is the relative contribution of modulation vs. block in the inhibition. Furthermore, kinetic modeling could allow estimation of the rate constants of the four major processes: access, egress, binding and unbinding.

### Simulations – Prototypical mechanisms

#### Model A. Non-state-dependent model

In the first model we simply allowed drug association to all four states with no state-dependent differences in accessibility (access/egress rates) or affinity (binding/unbinding rates). The egress constant (**K_E_**) was chosen to be 1000 μM, i.e., the rate of egress equaled the rate of access when the concentration was 1000 μM. In this simplest model we chose the access constant to be 10^-4^ ms^-1^μM^-1^, therefore at 1000 μM concentration bot access and egress proceeded with a rate constant of 0.1 ms^-1^. Binding and unbinding constants were both chosen to be 1 ms^-1^, i.e., neither access, nor binding was made energetically favorable, therefore at equilibrium channels were equally distributed between vacant, accessed (**AX**), and bound (**BX**) states (see the energy diagrams in Fig. 4A). In this, and subsequent models we show the results of simulations using the **SSI**, **RFI**, and **3PT** protocols. The effects on SSI and RFI protocols are always shown at 10, 100, and 1000 μM concentrations (Fig. 4A). The amplitude was decreased by channel block, but no shift of the **SSI** or **RFI** curves was observed (see insets in the **SSI** and **RFI** curves). The effect of the 260 ms inhibitor pulse (protocol **3PT**) was simulated at concentrations 1, 3.16, 10, 31.6, 100, 316, 1000, 3162, and 10000 μM. The concentration – inhibition curve (inset in the right panel of Fig. 4A) shows that the apparent affinity (500 μM) was lower than the egress constant (1000 μM), because of the coupled equilibria (vacant – accessed, and accessed – bound equilibria).

**Figure 4.**
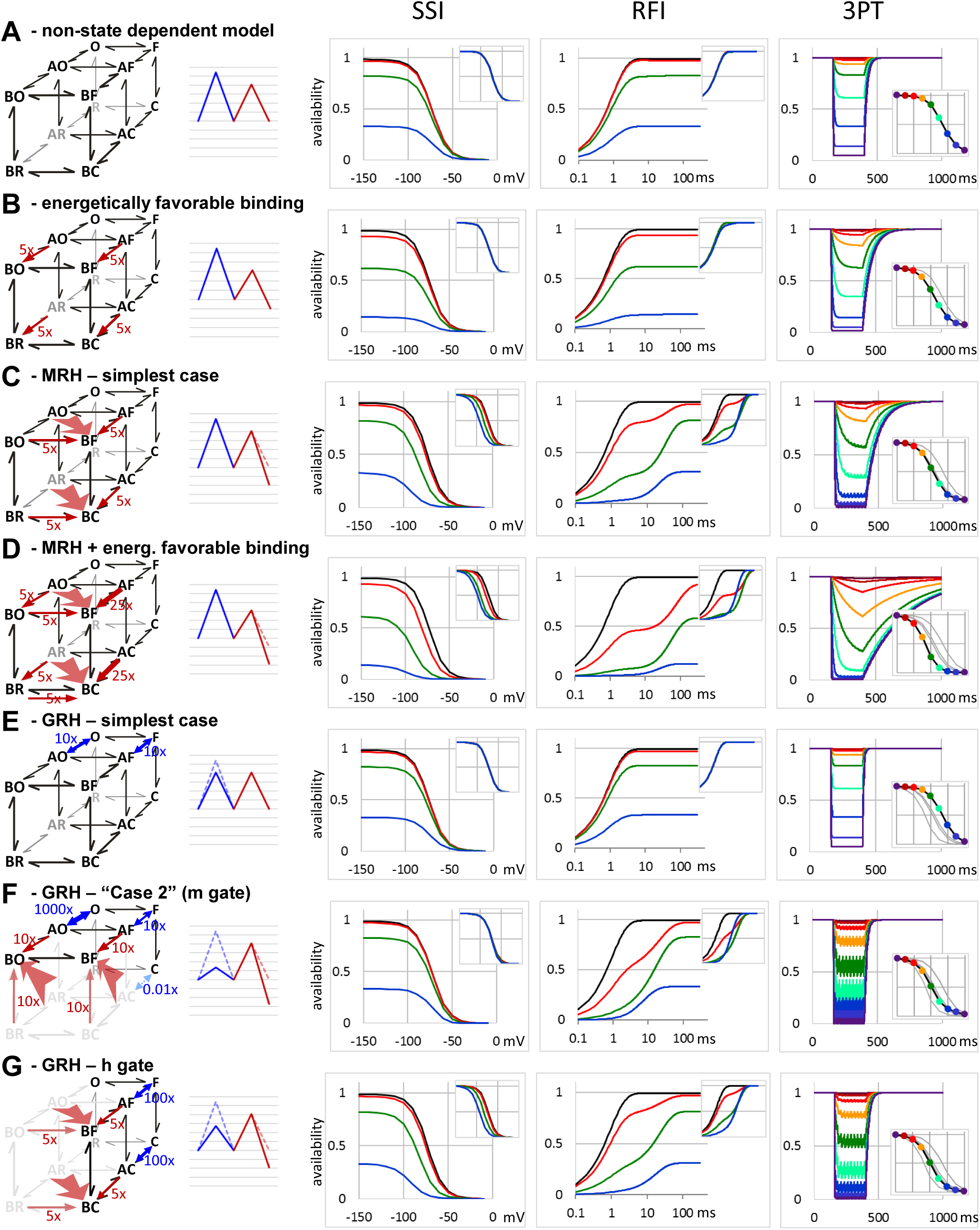
Simulations with prototypical mechanisms. In all cases 1^st^ column shows schematic mechanism of action, 2^nd^ column shows the energy diagram, 3^rd^, 4^th^ and 5^th^ columns the results of simulations in the **SSI**, **RFI,** and **3PT** protocols, respectively. Only vacant channels were considered available. Insets show normalized data for **SSI**, and **RFI** simulations, and concentration-inhibition curves calculated from the **3PT** simulation data. Colors in all simulations and the concentration-inhibition plot indicate concentrations 1 μM – dark purple, 3.16 μM – dark red, 10 μM – red, 31.6 μM – yellow, 100 μM – green, 316 μM – turquoise, 1000 μM – blue, 3160 μM – dark blue, 10000 μM – dark violet. In the schemes blue arrows indicate increased accessibility (decreased energy barrier), red arrows indicate increased affinity (in the examples of this figure binding rate was kept unchanged, affinity was increased by decreasing unbinding rate only). Wide transparent red arrows indicate allosteric factors, which modify both binding and gating equilibria. In the energy diagrams blue color indicates access/egress, red color indicates binding/unbinding. Pale, dashed lines indicate energy diagram for the R – AR – BR axis, continuous lines for the F – AF – BF axis for all rows except row (F), where it shows the O – AO – BO axis. In **SSI**, and **RFI** simulations only control and three concentrations are shown for clarity: 10, 100 and 1000 μM (red, green and blue lines, respectively). Only the first 1000 ms is shown in the **3PT** protocols. Concentration response curves always show data from previous models for comparison (thin gray lines). (A) non-state-dependent model. (B) Non-state-dependent model, with energetically favorable binding. (C) Simplest case of MRH. Fast inactivated state preference. (D) Fast inactivated case preference (MRH), with added energetically favorable binding. (E) Simplest case of GRH. Increased accessibility to channels with open activation gate. (F) “Case 2” GRH. The position of the activation gate determines accessibility. (G) GRH, the position of the inactivation gate determines accessibility.

#### Model B. Non-state-dependent model, with energetically favorable binding

It seemed more realistic if binding was made energetically favorable. In this model we simulated the case when binding was 5 times faster than unbinding, but binding and unbinding rates were not state-dependent. As we can see in Fig. 4B, the apparent affinity increased to 167 μM, but no shift of the **SSI** or **RFI** curves by the drug appeared.

#### Model C. Fast inactivated state preference, the simplest case of MRH

An allosteric factor (**AFB**) of 5 was introduced. This means that the rate of unbinding became 25-fold slower for inactivated channels (**BF → AF**, and **BC → AC**), and at the same time, the recovery from inactivation (**BF → BO,** and **BC → BR**) also became 25-fold slower (by default both **GFB** and **GFBg** were equal to 1/**AFB**, see Methods and Table S1). The leftward shift of the **SSI** curve, which is a characteristic property of most sodium channel inhibitors, was evident (Fig. 4C). Recovery curves became visibly biexponential at concentrations 10 and 100 μM, the first component representing recovery of vacant channels, while the second component representing both unbinding-followed-by-recovery and recovery-followed-by-unbinding processes. At 1000 μM concentration the recovery curve again became monoexponential, indicating close to full occupancy of the binding site within the first few milliseconds of the hyperpolarizing gap. The same concentration did not cause full inhibition in the **3PT** protocol, where depolarizations were not long enough to allow equilibration. The apparent affinity based on the **3PT** protocol was 127 μM.

#### Model D. Fast inactivated state preference, MRH, with added energetically favorable binding

Adding energetically favorable binding to the fast inactivated state preference further slowed both onset and offset, and increased apparent affinity in all protocols. While in itself favorable binding did not cause shift in the **SSI** protocol, it could intensify the shift caused by the MRH mechanism (Fig. 4D), and also further increased the apparent affinity calculated from simulations with the **3PT** protocol (45.2 μM).

#### Model E. GRH, Open state accessibility

The simplest case of GRH is when accessibility is increased depending on the position of one of the gates. We simulated increased accessibility for both open activation gate (*Model #5,* **GOA** = 10, Fig. 4E) and closed inactivation gate (**GFA** = 10, not shown). As compared to *Model#1,* onset and offset kinetics were accelerated in both cases, but there was no observable change in either concentration-inhibition curves, or **SSI** and **RFI** curves.

#### Model F. GRH, m gate, “Case 2”

In the original description of the GRH, Starmer, Grant and Strauss discussed three distinct models. The one named “case 2” is the most important one, because this was used to model the effects of lidocaine, etidocaine and QX compounds (Starmer et al., 1984). This model assumed that drug binding prevents closing of the activation (“m”) gate, therefore, in our current model, **AR**, **AC**, **BR** and **BC** states should not have significant occupancy (Fig. 4F, left panel). We implemented this mechanism by assigning a high guard factor (**GOA** = 1000) which accelerated access to- and egress from open activation gate states O and F. In addition, because accessibility to inactivated channels was supposed to be less than to open channels, we also assigned a guard factor to inactivated states: GFA = 0.01. The relative accessibility of **R, C**, **O**, and **F** states were thus 1, 0.01, 1000, and 10, respectively. This model (as well as other combinations of **GOA** and **GFA** within the 0.01 to 10000 range) changed onset and offset kinetics, but completely failed to change either the concentration response curve, or any of the **SSI** and **RFI** curves (not shown).

This was not unexpected, since we have not yet implemented the “hidden MRH element” (see section 2.1.1.), *i.e.,* the assumption that drug binding prevents closing of the activation gate. When we did so by introducing an allosteric factor (**AOB** = 10) that impeded closing of the activation gate of drug-bound states (**BO** and **BF**), the result was a significant increase in apparent affinity (110 μM), with a significant alteration of the **RFI** curve. The latter became clearly bi-exponential at 10 μM (but monoexponential again at 100 and 1000 μM where the binding sites were almost fully occupied). The time constant of the fast component (which corresponded to recovery of unbound channels) was unchanged, only its contribution to the amplitude decreased with increasing drug concentration. The second time constant reflected dissociation. In spite of the more than tenfold slowing of the recovery at concentrations 100 and 1000 μM, the normalized **SSI** curves were minimally affected (−0.07, −0.45 and - 1.33 mV shift at 10, 100 and 1000 μM, respectively). This indicates that setting the affinity of open-activation-gate-states higher does not effectively cause shifted **SSI** curves, therefore this mechanism alone could not account for the characteristic shift of these curves that is typical for most SCIs. We have experimented with different **GOA (**10 to 10 000), **GFA** (0.01 to 1) and **AOB** (1 to 25) values, as well as with making drug access voltage-dependent, but neither combination caused a significant shift of the **SSI c**urve. This, in fact, is in agreement with the results described in the original publication about the GRH, where for example the Kd of QX314 was set to 1.34 nM (!), while the shift of **SSI** curves (Figure 3 of (Starmer et al., 1984)) was illustrated at 1 mM concentration. If one calculates the shift of **SSI** curves using the exact formulas of the publication, an observable shift requires concentrations at least hundredfold higher than the K_d_. In contrast, in experiments most SCIs produce evident leftward shifts of the **SSI** curve around, and even below their K_d_ values (Lenkey et al., 2010, 2011).

#### Model G. GRH, h gate

Although at the time when the GRH was published the activation gate presented itself as an obvious physical correlate for the “guard”, recent structural and molecular dynamics data point to the fenestrations as further candidates for additional possible conformation-dependent “guards” (Boiteux et al., 2014; Yan et al., 2017). In other words, not only the hydrophilic-, but also the hydrophobic pathway may be guarded in a conformation-dependent manner. This suggests the GRH may be applicable not only to closed vs. activated channels, but also to non-inactivated vs. inactivated channels. It is important to investigate this possibility, as most SCIs can readily access inactivated channels even without activation. This raises the question whether increased apparent affinity to inactivated channels is due to increased accessibility, increased affinity, or both; and if both mechanisms are involved, what is their respective contribution. For this reason we also simulated the case when inactivated channels are more accessible. We assigned a high guard factor (**GFA** = 100) to closed-inactivation-gate-states. As we have discussed before (Model **E**), this in itself did not change apparent affinity and failed to modify **SSI** and **RFI** curves.

We then introduced a “hidden MRH element” that would cause inactivation (“h”) gate immobilization. Setting the **AFB** factor to 5 increased apparent affinity to 102.5 μM, a similar value to the model, described above. It also strongly shifted the **RFI** curve, and – most importantly – caused a shift of the **SSI** curves (-1.92, −10.0 and −20.4 mV shift at 10, 100 and 1000 μM, respectively).

In summary, the “pure” GRH (state-dependent accessibility without state-dependent affinity) was not able to evoke a shift of **SSI** curves, no matter which conformation was more accessible. The original “case 2” GRH model supposed m^3^ gate immobilization, which in fact is nothing else than modulation of gating by the drug. Even with this “hidden MRH element”, this model only caused minimal SSI shift, when it affected the open state (the m^3^ gate) only. This is due to the fact that the channel population spends relatively little time in open state. Only the GRH model with h gate immobilization *(i.e.* not only higher accessibility but also higher affinity to inactivated state, as suggested by the MRH) was able to reach an **SSI** shift that was comparable to experimentally observed values.

### Simulations – Quantitative investigation of specific hypotheses on inhibition mechanisms

#### Contribution of GRH and MRH to inhibition mechanisms

Thus far we have only shown individual examples for specific mechanisms. If one wants to be able to estimate the contribution of state-dependent affinity and state-dependent accessibility in the mechanism of inhibition of a drug, it is important to compare how different degrees of state-dependent accessibility and affinity are manifested in experimental results.

We started from the model with 10-fold increased accessibility and 10-fold increased affinity to inactivated states. First we investigated the effect of different affinities to inactivated states: we simulated 1-, 10-, and 100-times increased affinity (**AFB** = 1, 3.16, and 10). Then we investigated the effect of different accessibilities: we simulated 1-, 10-, and 100-times increased accessibility (**GFA** = 1, 10, and 100). In all cases we simulated the effect of 300 μM concentration, while affinity to resting state was set to 1000 μM, as in the previous section. The effects of increasing affinity and accessibility are compared in Fig. 5.

**Figure 5.**
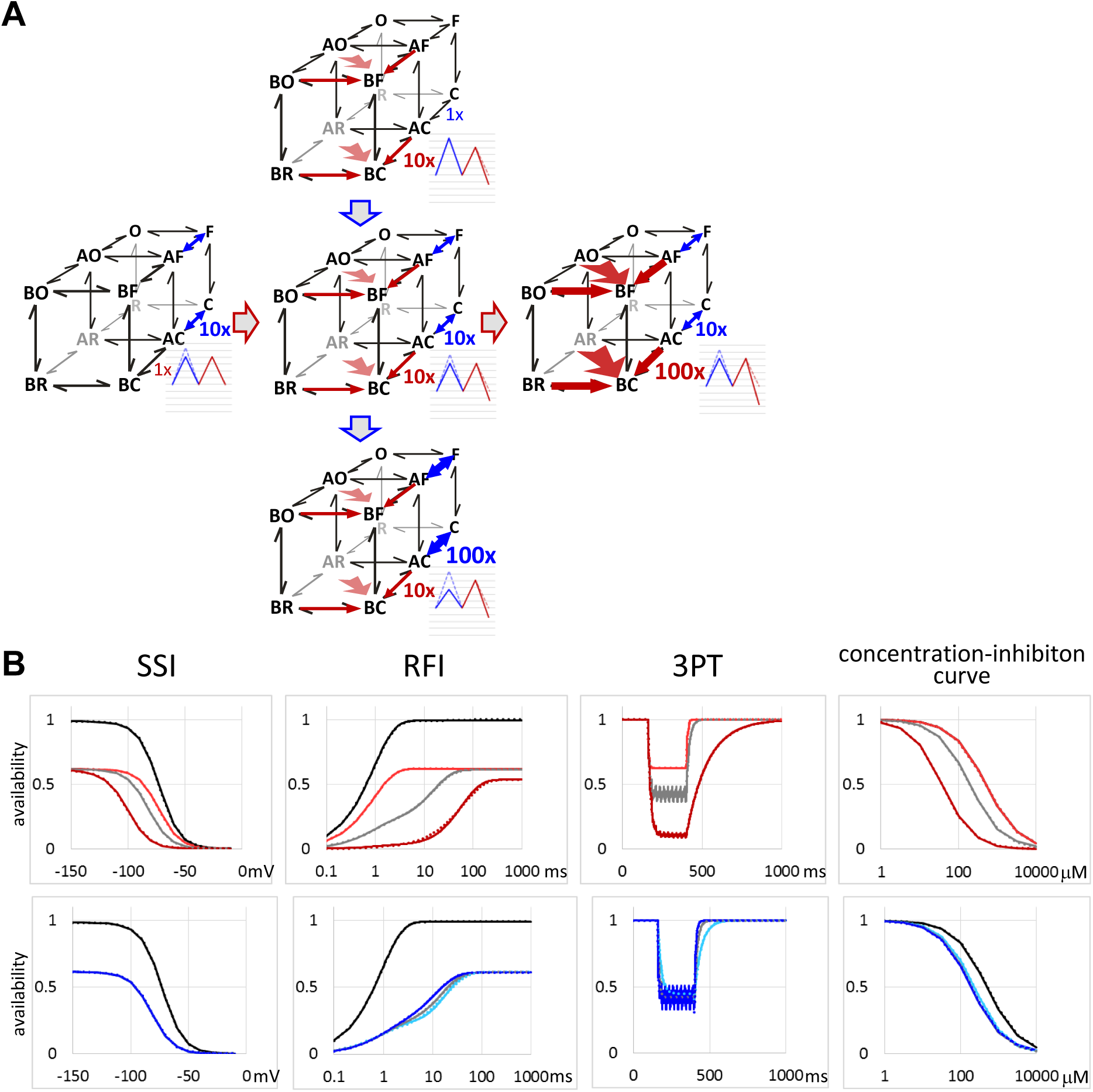
Simulation of different contributions of GRH and MRH to inhibition mechanisms. The effect of changing affinity vs. accessibility to inactivated state was investigated. (A) Schematic illustration of the five simulated parameter sets. For both affinity and accessibility to inactivated state, unchanged, 10-fold, and 100-fold increased cases were simulated. Width of arrows (blue for accessibility, red for affinity) and caption near arrows indicate the extent of state-dependence. In the energy diagrams blue color indicates access/egress, red color indicates binding/unbinding. Pale, dashed lines indicate energy diagram for the R – AR – BR axis, continuous lines for the F – AF – BF axis. (B) Results of the simulations. From left to right: **SSI**, **RFI**, **3PT** protocols, and concentration-inhibition curves from **3PT** protocol data. Black lines in the first two panels indicate control. Upper row: the effect of increased affinity. Light red, gray and dark red lines indicate 1x, 10x and 100x increased affinity, respectively. Lower row: the effect of increased accessibility. Light blue, gray and dark blue lines indicate 1x, 10x and 100x increased accessibility, respectively. Gray lines (the parameter set corresponding to the middle scheme) are identical in upper and lower panels. Results of fitting (see Methods) are shown by dotted line of the same color, overlaid on the simulated curves.

Increasing affinity to fast inactivated state in the **3PT** protocol caused more effective inhibition, with radically decelerated offset (8.5 ms, 18 ms, and 134 ms for **AFB** = 1, 3.16, and 10, respectively), and slightly decelerated onset (4.3 ms, 8.76 ms, and 15.9 ms). The apparent affinity (calculated from simulated concentration response curves) increased from 500 μM (**AFB** = 1) to 37.1 μM (**AFB** = 10). Resting state inhibition (to 61.7 % of control) was visible in all SSI curves, but the leftward shift was absent at **AFB** = 1. With increasing the allosteric factor, the SSI curve was progressively shifted, reaching −27.7 mV at **AFB** = 10. Similarly, at **AFB** = 1 the time constant of recovery from inactivation (0.925 ms) was almost indistinguishable from that of the control curve (0.918 ms). At **AFB** = 3.16 the fast component of the bi-exponential fit was still essentially unchanged (0.841 ms), but a slow component (τ = 15.9 ms) emerged, which contributed 71.0 % to the amplitude. At **AFB** = 10, the fast component almost completely disappeared, and the slow component had a time constant τ = 63.1 ms. The fast component reflected recovery of unbound channels, while the slow component reflected recovery of drug-bound channels, followed by unbinding and egress.

In contrast, increasing accessibility (while keeping increased affinity constant: **AFB** = 3.16) caused acceleration of onset and offset (τ_on_: 21.6, 8.76, and 4.21 ms; τ_off_: 41.9, 18.0, and 8.46 ms; for **GFA** = 1, 10, and 100, respectively), but only a mild increase in apparent affinity (from 230 μM at **GFA** = 1, to 192 μM at **GFA** = 100). It caused no shift at all in the SSI curve. Furthermore, not only did it not delay recovery from inactivation in the RFI protocol, but in fact accelerated it (from 18.6 ms at **GFA** = 1, to 11.2 ms at **GFA** = 100) by facilitating drug egress (which affected only the slow component of recovery).

We can conclude that the GRH mechanism can contribute somewhat to increased apparent affinity, but both the leftward shifted **SSI** curve, and the delayed **RFI** curve are sure signs of the presence of the MRH mechanism. The involvement of the GRH mechanism, on the other hand, is predominantly reflected in the acceleration of both onset and offset at depolarized membrane potentials.

#### Contribution of “lipophilic block” and “voltage sensor block”

Voltage sensor block has been defined as the component of inhibition that has the following properties: i) inhibitors act with high affinity, ii) the inhibition is voltage-dependent (because it requires open or inactivated conformation), iii) the inhibition is linked with stabilization of voltage sensors in domains III and IV (which can be seen in both the change of the gating charge vs. voltage (Q-V) relationship, and the shift of the **SSI** curve), and iv) it is dependent on the presence of the DIVS6 Phe (Hanck et al., 2009). More exactly, it is dependent on the aromatic π electron cloud of this residue. When the negative electrostatic potential of the aromatic ring has been partially or completely distorted by unnatural derivatives of the DIVS6 Phe, both the left shift of the **SSI** curve and the shift of the **RFI** curve were progressively suppressed and then eliminated, depending on the degree of distortion (Ahern et al., 2008). Knowing that the DIVS6 Phe residue is essential in mediating modulation of gating, we could simulate mutation of this residue by eliminating state preference (**AFB** = 1). We tested if elimination of state preference in simulations would have similar effect in simulations as introducing the F1579A mutation in experiments with riluzole and lidocaine; these results will be discussed below.

#### Contribution of channel block and modulation

It is generally assumed that bound SCI molecules inevitably cause full channel block. Our results with riluzole made us question this assumption, and forced us to suppose that SCI binding may cause inhibition predominantly by block, by modulation, or by both mechanisms. In the case of riluzole the predominant mechanism seems to be modulation without significant channel block. To explore the effect of incomplete block mechanism, we simulated different degrees of block. In order to do that we needed to redefine what we consider “available”. Thus far we defined available fraction as the ratio of vacant resting (**R**) channels, but now we redefined it as **R** + (1-**BlA)*AR** + (1-**BlB)*BR**, where **BlA** and **BlB** are blocked fraction of accessed and bound channels, respectively. For the sake of simplicity we chose to make **BlA** = **BlB**, and we will refer to their value as “**Bl**”. We have simulated different **Bl** values: 0, 0.25, 0.5, 0.75, and 1; for the sake of clarity we only show values: 0, 0.5, and 1 in Fig. 6. Because incomplete block decreased apparent affinity, we increased resting affinity to 100 μM. We simulated the case when both affinity and accessibility to inactivated channels are increased tenfold (**AFB** = 3.16, **GFA** = 10). The concentration of the simulated inhibitor was 100 μM (Fig. 6).

**Figure 6.**
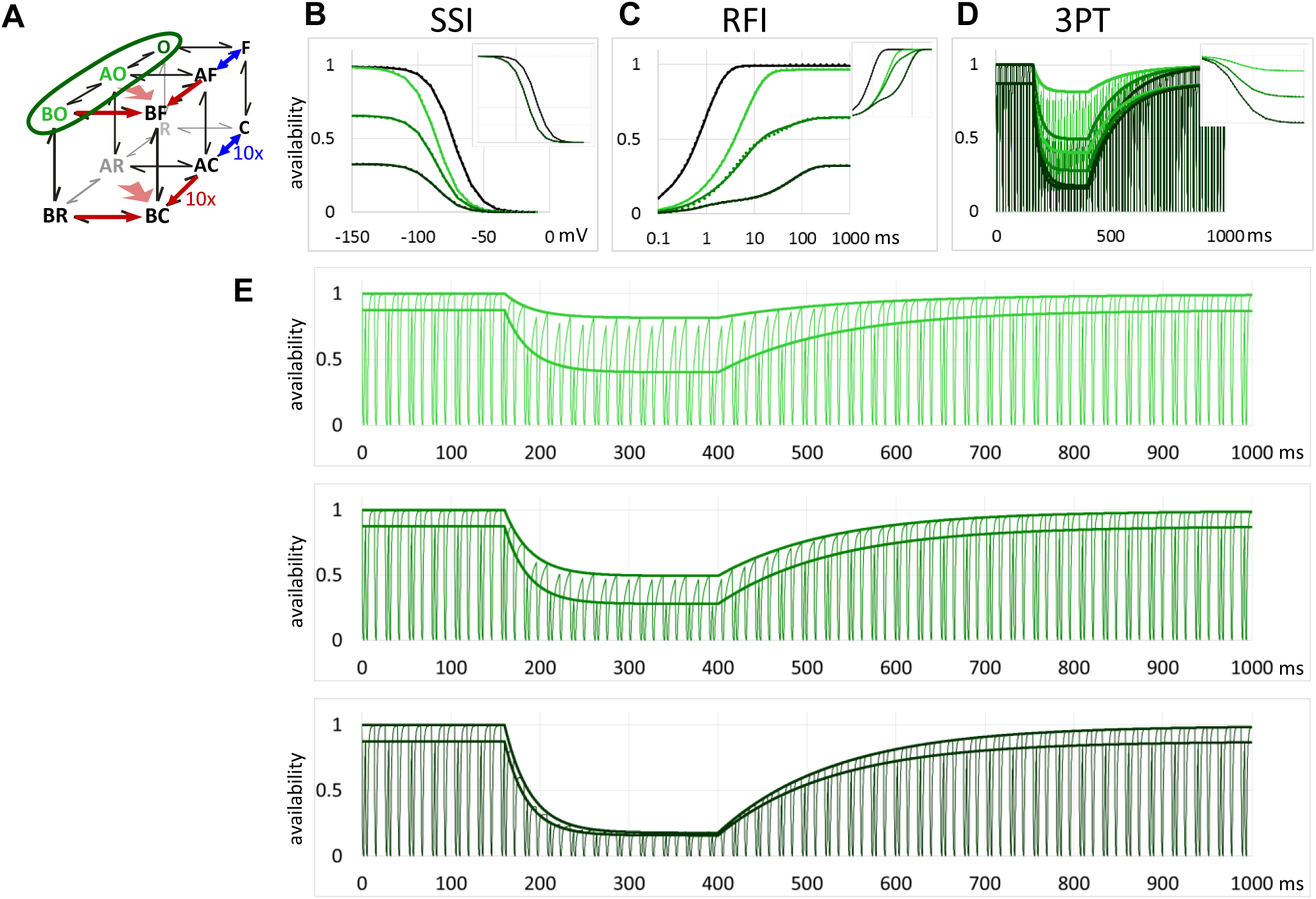
Simulation of non-blocking modulation. The simulations show what phenomena would be expected if accessed and/or bound open states (circled with green line in the scheme) were partially or fully conductive. Dark green traces show full block in accessed and bound states, *i.e.,* availability was defined as the fractional occupancy of R state alone. Medium green traces show partial block at accessed and bound open states; availability was calculated as R + 0.5 AR + 0.5 BR (where R, AR and BR indicate fractional occupancy of these states). Light green traces show non-blocking accessed and bound open states; availability was calculated as A + AR + BR. (A) Schematic illustration of the model. (B) **SSI** curves were normalized to the control amplitude, or in the inset, to their own maxima. (C) Shift of RFI curves, normalized to the control amplitude. Inset: curves normalized to their own maxima. (D) and (E) availability in the **3PT** protocols using full block (**bl** = 1), partial block (**bl** = 0.5), and no block (**bl** = 0) models. Thin lines indicate availability as it changes throughout the simulated experiment, thick lines are the linear (t < 130 ms), and exponential (130 ms < t 390 ms, and t > 390 ms) fits to peak availability values before the 1^st^ and the 2^nd^ pulses. (D) shows the three simulations together. (E) shows the three cases separately on an expanded time scale.

Making accessed and bound open states conducting did not shift the **SSI** curve, only the extent of resting inhibition depended on the value of **Bl.** However, decreasing **Bl** changed **RFI** curves radically. The control curve (without drug) recovered monoexponentially, with a time constant of 0.934 ms (when the exponential was on the first power). When we simulated the presence of 100 μM drug, recovery curves were either bi-, or tri-exponential. The fastest component contributed 7.2 to 10.8% of the control amplitude, and had a time constant between 0.78 and 0.91 ms, which reflected recovery of unbound channels (F → R). The intermediate component had a time constant between 6.06 and 6.27 ms, it contributed 86.2% of the control amplitude in the **Bl** = 0 model (light green in Fig. 6C), and 43.1% in the **Bl** = 0.5 model (medium green in Fig. 6C). It reflected modulated recovery of bound channels (BF → BR), therefore, it was present in all simulations, where channel block was not full (**Bl** < 1). The slow component had a time constant between 63.0 and 65.0 ms, it contributed 24.7% of the control amplitude at **Bl** = 1 (which was 76.5% of total recovery in this model – see dark green line in Fig. 6C), 13.0% at Bl = 0.5 (medium green in Fig. 6C), and it totally disappeared at **Bl** = 0 (light green in Fig. 6C). It reflected unbinding and egress. It is important to note that the RFI curve of the **Bl** = 0.5 model resembled the data obtained with 300 μM lidocaine, while the pattern of the **Bl** = 0 model was similar to data with riluzole, except that the two drugs probably had saturated binding sites, because no first component was observable. The **3PT** protocol showed the dynamic changes in availability throughout the experiment ((Fig. 6D and E). Peak availability values at the beginning of 1^st^ and 2^nd^ pulses were fitted separately. At the no-block case we saw the characteristic pattern observed at experiments with riluzole. The 2 to 10 ms hyperpolarized sections between depolarizing pulses were not long enough for unbinding and egress, but still allowed gating (recovery) of bound channels. Therefore, in the case of 1^st^ and 3^rd^ pulse-evoked currents the apparent inhibition much underestimated the actual binding. Incomplete block resulted in a divergence of amplitudes evoked by 1^st^ and 2^nd^ pulses, as we have seen in the case of riluzole. The inhibition, however, could still be correctly calculated from time constants of onset and offset. We can see that the non-blocking modulation could give a plausible explanation for the inhibition pattern seen with riluzole Could there be an alternative explanation for the divergence of 1^st^ and 2^nd^ pulse-evoked amplitudes, as well as for the shifted-but-monoexponential **RFI** curve? We have investigated this question, as it will be described in the next section.

In the case of the partial-block model (**Bl** = 0.5), we observed three subsequent processes: recovery of unbound channels, recovery of bound channels, and finally unbinding followed by egress. Time constants did not change considerably, only their ratio changed with the extent of block. In the case of lidocaine we saw a similar pattern, except that the first component of unbound channel recovery was missing. This indicates that recovery of drug-bound channels, followed by unbinding and egress of the drug, could explain the bi-exponential nature of **RFI** curves. Alternative explanations have also been examined, as we will discuss in the next section.

### Simulations – Understanding drug mechanisms

We wanted to see if we can propose a plausible hypothesis for the behavior of the two drugs, especially for the unique mode of action of riluzole. To accomplish this, we tried to reproduce experimentally found inhibition patterns in simulations. This allows us to give a quantitative estimation of the mechanism of inhibition, and in addition provides a way to test alternative hypotheses.

#### Riluzole

As we have discussed in the Introduction, the major elements of the peculiar inhibition pattern caused by riluzole are: i) a strong shift of the **SSI** curve with shallow resting inhibition; ii) peculiar shape of the **RFI** curve, which showed almost full inhibition at <1ms interpulse intervals, a steep loss of inhibition between 1 and 10 ms, and a shallow inhibition at >10 ms interpulse intervals; iii), the disagreement between the extent of inhibition and time constants of onset and offset in the **3PT** protocol.

We supposed, that onset and offset time constants reliably reflected occupancy of the binding site, from which it followed that the extent of inhibition did not, and a significant part of riluzole-bound channels must have conducted ions. One alternative explanation for the disagreement could be, to suppose two distinct binding sites for riluzole: the conventional blocking-and-modulatory site and an additional non-blocking-modulatory site. Simulations with two binding sites indeed could reproduce major characteristics of the inhibition (not shown). However, if there was a separate non-blocking-modulatory site, then mutation of the conventional binding site would leave this non-blocking modulator effect intact. Mutation of the binding site phenylalanine to alanine (F1579A) decreased both block and modulation (see experimental data above), and abolished the distinctive inhibition pattern of riluzole. This indicates that riluzole produces this special inhibition pattern by binding to the conventional local anesthetic binding site.

This leaves us with the first explanation, *i.e.,* the hypothesis that binding of riluzole to the conventional local anesthetic binding site does not prevent conduction by channel block mechanism, but predominantly by modulation. For this reason we simulated the case of incomplete channel block. Acceptable reproduction of the inhibition pattern required ~500- to 4000-fold increased affinity of inactivated channels (5.5 < **AFB** < 8, and 4 < **AOB** < 8; increase in affinity to inactivated channels is calculated as **AFB**^2^ * **AOB**^2^), with a rather low increase of accessibility (both **GFA** and **GOA** < 10, while their product was between 25 and 50). In addition, and most importantly, block values for bound channels (BlB) had to be below 0.05 in order to reproduce the steep **RFI** curve, as well as the different extent of inhibition by the 1^st^ and 2^nd^ pulses of the **3PT** protocol. Simulations with one set of parameters (see Table S1) are shown in Fig. 7A to F. In summary, to produce an inhibition pattern similar to the effect of riluzole, we needed to introduce moderately increased accessibility and strongly increased affinity to inactivated state, and we needed to suppose that drug-bound open channels can conduct almost as effectively as vacant open channels (conductance of accessed and bound channels is at least 95% of the conductance of vacant channels).

**Figure 7.**
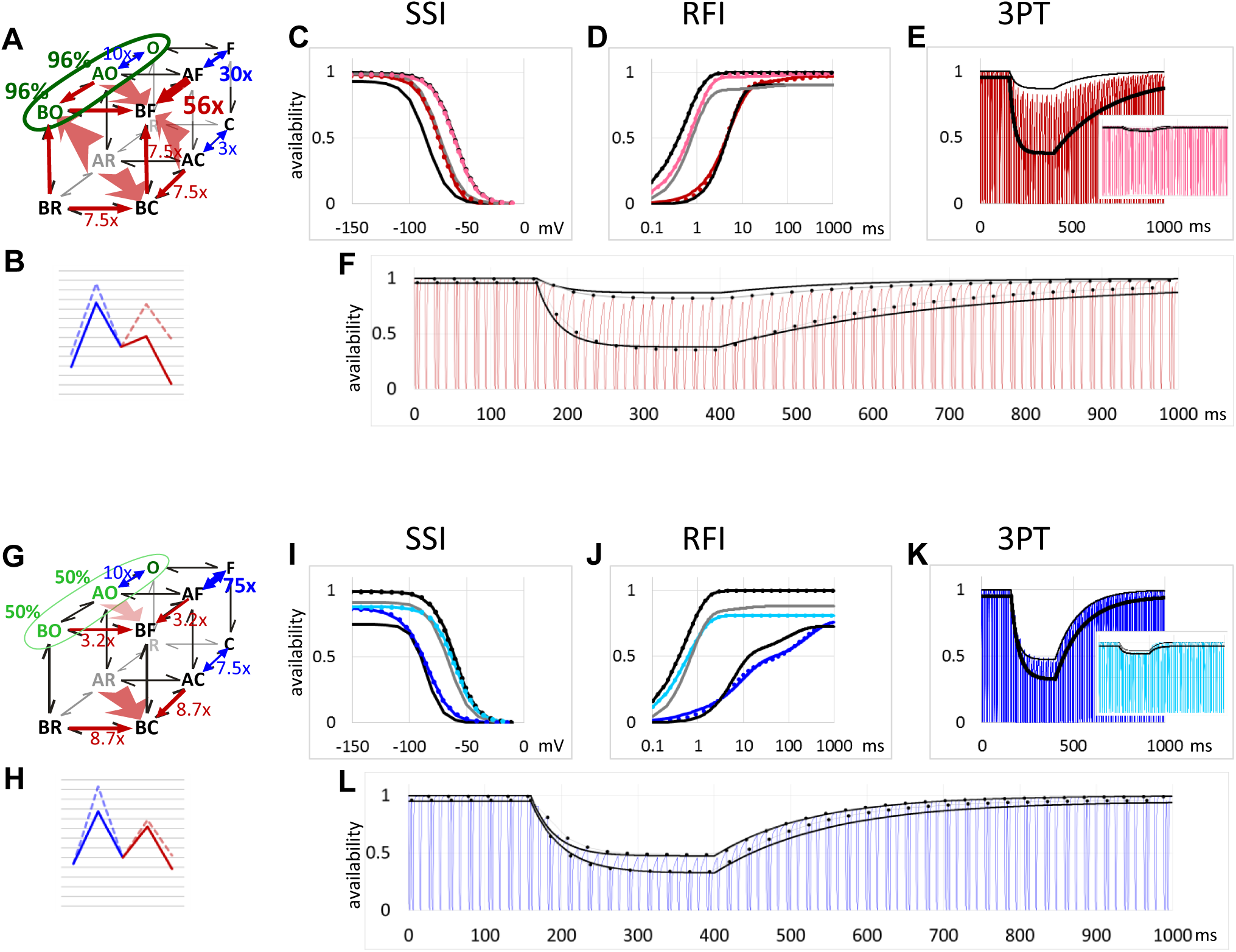
Simulation of the effect of riluzole and lidocaine in WT and mutant channels. The attempt to reproduce the results with riluzole and lidocaine is shown in panels A to F, and G to L, respectively. Red and pink lines in panels A to F indicate simulation of riluzole effects on wild type and on F1579A mutants, respectively. Dark blue and light blue lines in panels G to L indicate simulation of lidocaine effects on wild type and on F1579A mutants, respectively. (A) and (G) Schematic illustration of the proposed mode of action of riluzole and lidocaine. Width of arrows (blue for accessibility, red for affinity) and caption near arrows indicate the extent of state-dependence. Wide transparent red arrows indicate allosteric factors. Green fonts indicate conducting states, caption indicate percentage of vacant channel conductance. (B) and (H) Energy diagrams for the model. Blue color indicates access/egress, red color indicates binding/unbinding. Pale, dashed lines indicate energy diagram for the R – AR – BR axis, continuous lines for the F – AF – BF axis. (C) and (I) Simulated **SSI** curves, normalized to the control amplitude. Dotted lines indicate Boltzmann fits. Black and gray lines show the mean of experimental data from wild type and F1579A mutant channels, respectively. (D) and (J) Simulated **RFI** curves, normalized to the control amplitude. Dotted lines indicate exponential fits. Black and gray lines show experimental data on WT and mutant channels, respectively. (E) and (K) Simulated availability in the **3PT** protocols. The mean of curves fit on the normalized peak amplitudes of 1^st^ and 2^nd^ pulse-evoked currents are shown by thin and thick black lines, respectively. Insets show data on mutant channels, together with simulations of mutant channels (AFB = 1). (F) and (L) The same data as in panels E and K, on an expanded time scale. Peak availability values of simulations at the beginning of 1^st^ and 2^nd^ pulses are shown by dotted lines to help comparing experimental and simulated data.

We also attempted to simulate the alternative explanation, by supposing full block for accessed and bound channels, and reproducing the **RFI** curve by setting unbinding and egress sufficiently fast. Fig. 8 shows the results of the simulation. The essential difference is best seen on the energy diagrams (compare Fig. 7B and Fig. 8B). To reproduce the RFI curve, egress rates had to be increased 27.4-fold for the AR and 86.6-fold for the AF → F transition. The shift of the **RFI** curve was adequately reproduced, and the shift in the **SSI** curve was similar to the one we got with the model shown in Fig. 7. However, because access and egress had to be set very fast, this model completely failed to reproduce the slow offset seen in **3PT** protocols; the offset was too fast to resolve, there was full recovery from one train to the next one. We can conclude that both alternative hypotheses failed to explain the inhibition pattern produced by riluzole.

**Figure 8.**
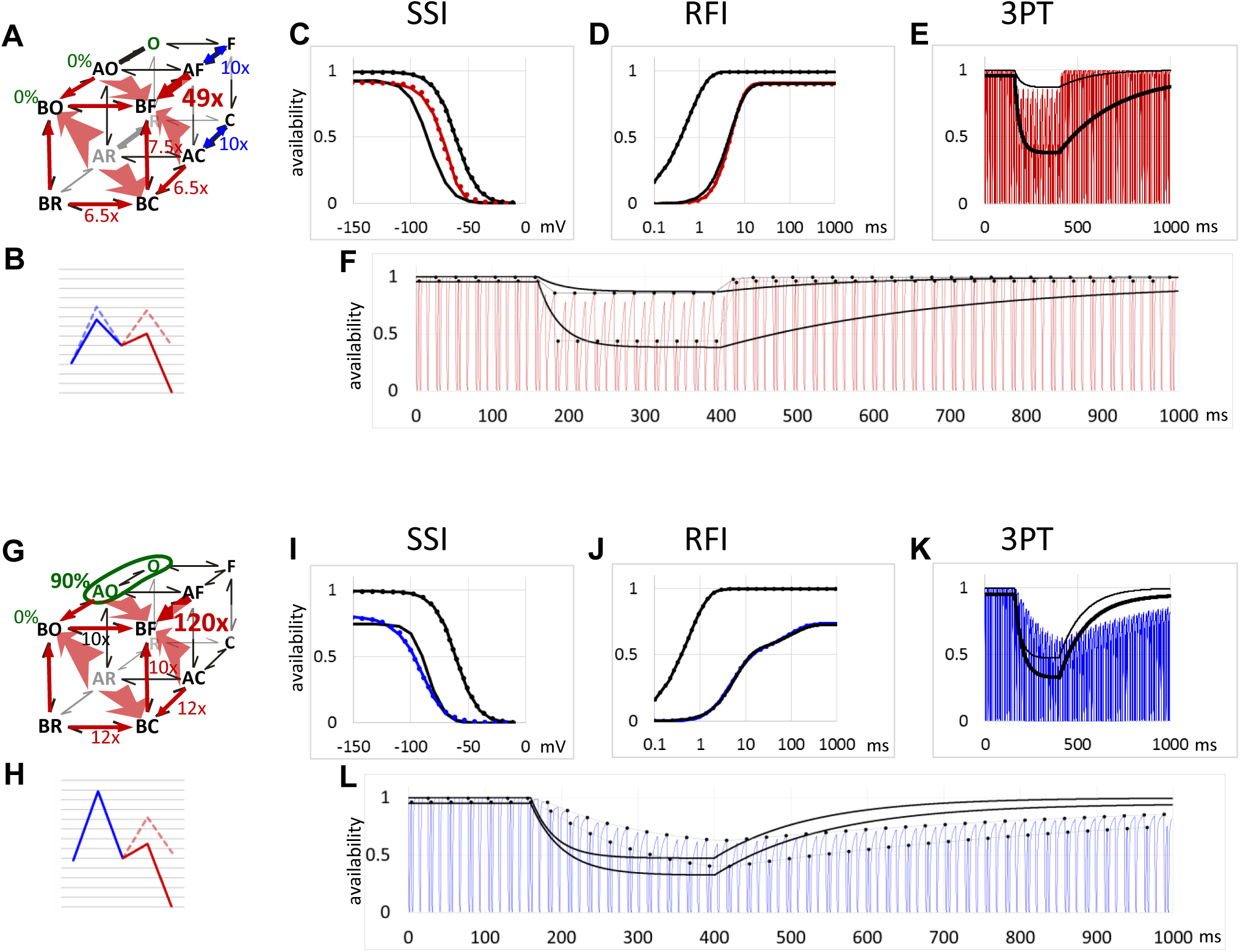
Simulation of the effect of riluzole and lidocaine supposing full block by bound inhibitors. Simulations made to reproduce experimental data with riluzole and lidocaine are shown in panels A to F, and G to L, respectively. Red and blue continuous lines show simulation of the effects of riluzole and lidocaine, respectively. (A) and (G) Schematic illustration of the alternative model for riluzole and lidocaine. Width of arrows (blue for accessibility, red for affinity) and caption near arrows indicate the extent of state-dependence. Wide transparent red arrows indicate allosteric factors. Green fonts indicate conducting states, caption indicate percentage of vacant channel conductance. (B) and (H) Energy diagrams for the model. Blue color indicates access/egress, red color indicates binding/unbinding. Pale, dashed lines indicate energy diagram for the R – AR – BR axis, continuous lines for the F – AF – BF axis. (C) and (I) Simulated **SSI** curves, normalized to the control amplitude. Dotted lines indicate Boltzmann fits. Black lines show experimental data. (D) and (J) Simulated **RFI** curves, normalized to the control amplitude. Dotted lines indicate exponential fits. Black lines show experimental data. (E) and (K) Simulated availability in the **3PT** protocols. The mean of curves fit on the normalized peak amplitudes of 1^st^ and 2^nd^ pulse-evoked currents are shown by thin and thick black lines, respectively. (F) and (L) The same data as in panels E and K, on an expanded time scale. Peak availability values of simulations at the beginning of 1^st^ and 2^nd^ pulses are shown by dotted lines to help comparing experimental and simulated data.

Based on the data in literature we expect that the aromatic ring of F1579 forms an electrostatic interaction with the ligand, which is responsible for the major part of the binding energy. Therefore, the effect of mutation could be simulated by eliminating preferential affinity to inactivated state (**AFB** = 1), while leaving all the rest of the parameters unchanged. When we simulated this case, inhibition patterns resembled those obtained in mutant channels (Fig. 7), but, interestingly, the model underestimated the shift of **SSI** curves (similarly to simulation of WT channels). In mutant experiments **SSI** curves were shifted by −12.1 mV, which was not reproduced in simulations. The reason why riluzole caused an unexpectedly large shift of **SSI** curves in both WT and mutant channels needs to be further studied, it might indicate binding to an additional binding site, in which F1579 is not involved.

#### Lidocaine

We experimented with the parameters to see what was required to reproduce the inhibition pattern of lidocaine. Reproducing the extent of shift (-24.2 ± 3.96 mV in experiments) of **SSI** curves required **AFB** to be between 4 and 9. However, unlike in the case of riluzole, **AOB** (which only minimally affected V_1/2_ of the **SSI** curve) could not be higher than unity, it was between 0.3 and 0.8 in the best performing parameter sets.

For the fast component of **RFI** curves, the extent of shift was predominantly determined by **AFB** and **GFBg** (guard factor of inactivation). We checked the range of reasonable **GFBg** values (between 1/**AFB** and 1), and found that **AFB** values had to be between 3 and 9 in order to reproduce the ~8-fold prolongation of the first component of recovery. We can conclude that data from SSI and RFI protocols agree that **AFB** must be between 4 and 9.

As we have mentioned, the **RFI** curve in the presence of 300 μM lidocaine was biphasic, the fast component (4.09 ± 0.53 ms) contributed ~70 % of the amplitude, and the slow component (103.5 ± 12.4 ms) the remaining ~30 %. We aimed to understand which exact processes constituted the two components. One interpretation was that the fast component represented modulated recovery from inactivated state, and the second component unbinding and egress. In this case we needed to suppose that lidocaine-bound channels have a certain (limited) conductance. Setting the block parameter to 0.3 < **Bl** < 0.6 together with 4 < **AFB** < 9, and a strong state-dependent accessibility (50 < **GFA**\***GOA** < 500) could acceptably reproduce the inhibition pattern produced by lidocaine in all three protocols. While in the case of riluzole both **AFB** and **AOB** contributed to state-dependent affinity, in the case of lidocaine we found that **AOB** had to be ≤ 1. Simulations with one set of parameters (see Table S1) are shown in Fig. 7G to L.

Another interpretation could be that the fast component of the **RFI** curve reflected the parallel processes of delayed-recovery-followed-by-rapid-unbinding and delayed-unbinding-followed-by-rapid-recovery; and the slow component reflected drug egress. In this case we must suppose that although lidocaine-bound channels cannot conduct, lidocaine-accessed channels can, and this limited conductance forms the plateau between the first and second phases of the **RFI** curve. We could reproduce the experimental **RFI** curve very accurately by setting **BlB** = 1 (i.e., lidocaine-bound channels were fully blocked), but allowing lidocaine-accessed channels to conduct (0 < **BlA** < 0.25) (Fig. 8J). In this case, we needed to increase modulation, and to slow down egress (compare Fig. 7H and Fig. 8H). This high energy barrier in the access/egress step, however, was irreconcilable with onset and offset rates in the **3PT** protocol: onset and especially offset became much slower than in experiments (Fig. 8K and L).

Investigation of F1579A mutant channels may also help to distinguish between alternative interpretations. One would expect that the mutation causes unbinding to be accelerated, while egress, or partitioning should not be affected. We observed that this single mutation totally abolished the slow component of recovery (which was within the 73.9 to 155 ms range), and also radically accelerated the fast component. This result strongly suggests that the fast component of **RFI** curves in wild type channels indeed reflected recovery of lidocaine-bound channels, while the slow component reflected unbinding and egress.

In summary, simulations suggested that the mechanism of action for lidocaine is a definite modulation of gating, which explains the ~20 mV leftward shift of the **SSI** curve, and the ~8-fold decelerated recovery from inactivated state. This accounts for the first phase of recovery, which proceeds with a time constant of ~4 ms. Unbinding and egress constitutes the second phase, which proceeds with a time constant of ~150 ms. This implies that drug-bound channels must be able to conduct current, although with decreased conductance. Simulations with partial block could reproduce experimental data when lidocaine-bound channels could conduct 30 to 60 % of the total current. State-dependent accessibility had to be equal or even larger in magnitude than state-dependent affinity, suggesting that both MRH and GRH mechanisms contribute to the inhibition.

Based on the results of simulations we propose the following hypotheses regarding the mode of action of riluzole and lidocaine: Riluzole-bound channels can open and conduct ions, the channel block caused by the presence of riluzole at the binding site is negligible (less than 5% of riluzole-bound channels is blocked). In the case of lidocaine we estimate that 40 to 70 % of lidocaine-bound channels are blocked at a time. In the case of riluzole, state-dependent affinity is dominant (500 to 4000-fold increased affinity), and state-dependent accessibility plays only a minor role (25 to 50-fold increase). In contrast, state-dependent accessibility and affinity are equally important elements of the mechanism of action for lidocaine.

## DISCUSSION

Sodium channels are promising targets in different membrane excitability diseases, such as heart arrhythmias, muscle disorders, pain syndromes, epilepsy as well as several other neurological and psychiatric disorders. However, the family of sodium channels proved to be a difficult target, not because it is difficult to find compounds that exert an inhibitory effect, but because potent SCIs do not necessarily make potent drugs, and therefore looking for potent SCIs may not be the best strategy. It is still not clear, however, what exactly to look for. High state-dependence has been proposed to be an indicator of therapeutic usefulness (Cerne et al., 2016; Liu et al., 2011), but state-dependence is a property that is difficult to standardize, because it depends on several different properties of the experimental protocol. We propose that modulation is a preferable mechanism as compared to channel block, because it can be more selective to pathological activity patterns.

The important question is, how to tell apart modulation and channel block. Combined voltage- and drug application-protocols, like the one we called **3PT** can be used to monitor both modulated gating (within-train recovery) and genuine dissociation (between-trains recovery). In the case of “extreme” modulators, such as riluzole, where channel block seems to have very little contribution to the inhibition, the discrepancy between the rates of within-train and between-trains recovery evidently shows that non-blocking modulation must exist and must be responsible for the delay in RFI curves. In the case of less obvious modulators, like lidocaine, the biphasic nature of the **RFI** curve may give a hint. According to the conventional interpretation the **RFI** curve in the presence of the drug reflects dissociation in parallel with delayed (modulated) gating. When we attempted to simulate the effect of riluzole and lidocaine we found that when parameters were set according to this conventional interpretation (recovery within the ~1 to 10 ms range was caused by dissociation), then results of the **3PT** protocol could not be reproduced (Fig. 8).

In summary, we believe that the contribution of channel block and modulation can be assessed by a protocol where rapid drug application is combined with short trains, which monitor recovery from inactivation.

Our experiments with the F1579A binding site mutant identifies this residue as the most important residue that couples binding with modulation. In this respect it is both key to the MRH and to “voltage sensor block”, which is essentially an inhibition with a strong modulation component. The contribution of block and modulation can vary between SCI drugs, relatively strong block with weak modulation would give an inhibition similar to “lipophilic block”.

Non-blocking modulation may also be related to the phenomenon of rapid block. It is conceivable that for some drugs drug-accessed state produces a reduced conductance, such as during rapid block, and drug-bound state produces discrete block. However, as we have seen in the example of riluzole and lidocaine, individual drugs can behave widely differently, and no general conclusion can be drawn for SCIs.

As for the contribution of the GRH and the MRH, our main conclusion came from the study of prototypical mechanisms. We suppose that most real SCI mechanisms include both hypotheses, and they are certainly not mutually exclusive. We found that the presence of GRH is predominantly shown by accelerated onset and offset at depolarized membrane potentials. A shift in the SSI curve, on the other hand, is a definite proof that modulation contributes to the mechanism of inhibition. State-dependent accessibility alone could not cause a shift, and in the original description of GRH (“Case 2”) increased open state accessibility could only cause a minimal shift because modulation was also assumed.

Kinetic modeling provides a way to assess the relative contribution of GRH (state-dependent accessibility) and MRH (state-dependent affinity) in the mode of action of individual SCI drugs. We conclude that for riluzole and lidocaine both mechanisms contribute to state-dependent inhibition, but while in the case of riluzole state-dependent affinity, (and therefore modulation) are dominant, in the case of lidocaine state-dependent accessibility and channel block also significantly contribute to the inhibition.

## Abbreviations used

SCI: sodium channel inhibitor
MRH: modulated receptor hypothesis
GRH: guarded receptor hypothesis
SSI: steady-state inactivation protocol
RFI: recovery from inactivation
3PT: three pulse train
R, C, O, F, AR, AC, AO, AF, BR, BC, BO, BF: states of the model:
R: resting
C: closed inactivated
O: open
F: inactivated (fast inactivated)
AX: any of the accessed states BX – any of the bound states
AFB: allosteric factor stabilizing drug-bound fast inactivated state
AOB: allosteric factor stabilizing drug-bound open state
GFA: “guard factor”, accelerating access/egress to/from fast inactivated states
GOA: “guard factor”, accelerating access/egress to/from open states
GFBg: “guard factor for fast inactivation/recovery
BlA: blocked fraction for accessed channels
BlB: blocked fraction for bound channels
Bl: blocked fraction for accessed and bound channels
RIL: riluzole
LID: lidocaine

## ACKNOWLEDGMENTS

This work was supported by the Hungarian Brain Research Program (KTIA-NAP-13-2-2014-002). Plasmid DNA for WT and mutant NaV1.4 channel were kindly provided by Hannes Todt, pCaggs IgG-Fc plasmid DNA by Dr. Emilio Casanova. The authors declare no competing financial interests.

## REFERENCES

Ahern, C. A., Eastwood, A. L., Dougherty, D. A., and Horn, R. (2008). Electrostatic contributions of aromatic residues in the local anesthetic receptor of voltage-gated sodium channels. Circ. Res. 102, 86–94. doi:10.1161/CIRCRESAHA.107.160663.

Arcisio-Miranda, M., Muroi, Y., Chowdhury, S., and Chanda, B. (2010). Molecular mechanism of allosteric modification of voltage-dependent sodium channels by local anesthetics. J. Gen. Physiol. 136, 541–554. doi:10.1085/jgp.201010438.

Bagneris, C., Naylor, C. E., McCusker, E. C., and Wallace, B. A. (2014). Structural model of the open-closed-inactivated cycle of prokaryotic voltage-gated sodium channels. J. Gen. Physiol. 145, 5–16. doi:10.1085/jgp.201411242.

Bean, B. P., Cohen, C. J., and Tsien, R. W. (1983). Lidocaine block of cardiac sodium channels. J. Gen. Physiol. 81, 613–642.

Benoit, E., and Escande, D. (1991). Riluzole specifically blocks inactivated Na channels in myelinated nerve fibre. Pflüg. Arch. Eur. J. Physiol. 419, 603–609.

Boiteux, C., Vorobyov, I., and Allen, T. W. (2014). Ion conduction and conformational flexibility of a bacterial voltage-gated sodium channel. Proc. Natl. Acad. Sci. 111, 3454–3459.

Catterall, W. A., and Swanson, T. M. (2015). Structural Basis for Pharmacology of Voltage-Gated Sodium and Calcium Channels. Mol. Pharmacol. doi:10.1124/mol.114.097659.

Centonze, D., Calabresi, P., Pisani, A., Marinelli, S., Marfia, G. A., and Bernardi, G. (1998). Electrophysiology of the neuroprotective agent riluzole on striatal spiny neurons. Neuropharmacology 37, 1063–1070.

Cerne, R., Wakulchik, M., Krambis, M. J., Burris, K. D., and Priest, B. T. (2016). IonWorks Barracuda Assay for Assessment of State-Dependent Sodium Channel Modulators. Assay Drug Dev. Technol. 14, 84–92. doi:10.1089/adt.2015.677.

Cho, J.-H., Choi, I.-S., Lee, S.-H., Lee, M.-G., and Jang, I.-S. (2015). Contribution of persistent sodium currents to the excitability of tonic firing substantia gelatinosa neurons of the rat. Neurosci. Lett. 591, 192–196. doi:10.1016/j.neulet.2015.02.039.

Cifra, A., Mazzone, G. L., and Nistri, A. (2013). Riluzole: what it does to spinal and brainstem neurons and how it does it. Neurosci. Rev. J. Bringing Neurobiol. Neurol. Psychiatry 19, 137–144. doi:10.1177/1073858412444932.

Clairfeuille, T., Xu, H., Koth, C. M., and Payandeh, J. (2016). Voltage-gated sodium channels viewed through a structural biology lens. Curr. Opin. Struct. Biol. 45, 74–84. doi:10.1016/j.sbi.2016.11.022.

Del Negro, C. A., Morgado-Valle, C., and Feldman, J. L. (2002). Respiratory rhythm: an emergent network property? Neuron 34, 821–830.

Desaphy, J.-F., Carbonara, R., Costanza, T., and Conte Camerino, D. (2014). Preclinical evaluation of marketed sodium channel blockers in a rat model of myotonia discloses promising antimyotonic drugs. Exp. Neurol. 255, 96–102. doi:10.1016/j.expneurol.2014.02.023.

Eijkelkamp, N., Linley, J. E., Baker, M. D., Minett, M. S., Cregg, R., Werdehausen, R., et al. (2012). Neurological perspectives on voltage-gated sodium channels. Brain 135, 2585–2612. doi:10.1093/brain/aws225.

England, S., and de Groot, M. J. (2009). Subtype-selective targeting of voltage-gated sodium channels. Br. J. Pharmacol. 158, 1413–1425. doi:10.1111/j.1476-5381.2009.00437.x.

Fehlings, M. G., Nakashima, H., Nagoshi, N., Chow, D. S. L., Grossman, R. G., and Kopjar, B. (2016). Rationale, design and critical end points for the Riluzole in Acute Spinal Cord Injury Study (RISCIS): a randomized, double-blinded, placebo-controlled parallel multi-center trial. Spinal Cord 54, 8–15. doi:10.1038/sc.2015.95.

Fischer, B. D., Ho, C., Kuzin, I., Bottaro, A., and O’Leary, M. E. (2017). Chronic exposure to tumor necrosis factor in vivo induces hyperalgesia, upregulates sodium channel gene expression and alters the cellular electrophysiology of dorsal root ganglion neurons. Neurosci. Lett. 653, 195–201. doi:10.1016/j.neulet.2017.05.004.

Gawali, V. S., Lukacs, P., Cervenka, R., Koenig, X., Rubi, L., Hilber, K., et al. (2015). Mechanism of Modification, by Lidocaine, of Fast and Slow Recovery from Inactivation of Voltage-Gated Na^+^ Channels. Mol. Pharmacol. 88, 866–879. doi:10.1124/mol.115.099580.

Gingrich, K. J., Beardsley, D., and Yue, D. T. (1993). Ultra-deep blockade of Na+ channels by a quaternary ammonium ion: catalysis by a transition-intermediate state? J. Physiol. 471, 319–341.

Hanck, D. A., Makielski, J. C., and Sheets, M. F. (1994). Kinetic effects of quaternary lidocaine block of cardiac sodium channels: a gating current study. J. Gen. Physiol. 103, 19–43.

Hanck, D. A., Nikitina, E., McNulty, M. M., Fozzard, H. A., Lipkind, G. M., and Sheets, M. F. (2009). Using Lidocaine and Benzocaine to Link Sodium Channel Molecular Conformations to State-Dependent Antiarrhythmic Drug Affinity. Circ. Res. 105, 492–499. doi:10.1161/CIRCRESAHA.109.198572.

Hebert, T., Drapeau, P., Pradier, L., and Dunn, R. J. (1994). Block of the rat brain IIA sodium channel alpha subunit by the neuroprotective drug riluzole. Mol. Pharmacol. 45, 1055–1060.

Hille, B. (1977). Local anesthetics: hydrophilic and hydrophobic pathways for the drug-receptor reaction. J. Gen. Physiol. 69, 497–515.

Hondeghem, L. M., and Katzung, B. G. (1977). Time- and voltage-dependent interactions of antiarrhythmic drugs with cardiac sodium channels. Biochim. Biophys. Acta 472, 373–398.

Huang, C.-J., Harootunian, A., Maher, M. P., Quan, C., Raj, C. D., McCormack, K., et al. (2006). Characterization of voltage-gated sodium-channel blockers by electrical stimulation and fluorescence detection of membrane potential. Nat. Biotechnol. 24, 439–446. doi:10.1038/nbt1194.

Karoly, R., Lenkey, N., Juhasz, A. O., Vizi, E. S., and Mike, A. (2010). Fast- or slow-inactivated state preference of Na+ channel inhibitors: a simulation and experimental study. PLoS Comput. Biol. 6, e1000818. doi:10.1371/journal.pcbi.1000818.

Kimbrough, J. T., and Gingrich, K. J. (2000). Quaternary ammonium block of mutant Na+ channels lacking inactivation: features of a transition-intermediate mechanism. J. Physiol. 529 Pt 1, 93–106.

Koizumi, H., and Smith, J. C. (2008). Persistent Na+ and K+-dominated leak currents contribute to respiratory rhythm generation in the pre-Bötzinger complex in vitro. J. Neurosci. Off. J. Soc. Neurosci. 28, 1773–1785. doi:10.1523/JNEUR0SCI.3916-07.2008.

Kuo, J. J., Lee, R. H., Zhang, L., and Heckman, C. J. (2006). Essential role of the persistent sodium current in spike initiation during slowly rising inputs in mouse spinal neurones. J. Physiol. 574, 819–834. doi:10.1113/jphysiol.2006.107094.

Lenkey, N., Karoly, R., Epresi, N., Vizi, E., and Mike, A. (2011). Binding of sodium channel inhibitors to hyperpolarized and depolarized conformations of the channel. Neuropharmacology 60, 191–200. doi:10.1016/j.neuropharm.2010.08.005.

Lenkey, N., Karoly, R., Lukacs, P., Vizi, E. S., Sunesen, M., Fodor, L., et al. (2010). Classification of Drugs Based on Properties of Sodium Channel Inhibition: A Comparative Automated Patch-Clamp Study. PLoS ONE 5, e15568. doi:10.1371/journal.pone.0015568.

Liu, H., Tateyama, M., Clancy, C. E., Abriel, H., and Kass, R. S. (2002). Channel openings are necessary but not sufficient for use-dependent block of cardiac Na+ channels by flecainide evidence from the analysis of disease-linked mutations. J. Gen. Physiol. 120, 39–51.

Liu, Y., Beck, E. J., and Flores, C. M. (2011). Validation of a patch clamp screening protocol that simultaneously measures compound activity in multiple states of the voltage-gated sodium channel Nav1.2. Assay Drug Dev. Technol. 9, 628–634. doi:10.1089/adt.2011.0375.

Lounkine, E., Keiser, M. J., Whitebread, S., Mikhailov, D., Hamon, J., Jenkins, J. L., et al. (2012). Large-scale prediction and testing of drug activity on side-effect targets. Nature 486, 361–367. doi:10.1038/nature11159.

Lukacs, P., Gawali, V. S., Cervenka, R., Ke, S., Koenig, X., Rubi, L., et al. (2014). Exploring the structure of the voltage-gated Na+ channel by an engineered drug access pathway to the receptor site for local anesthetics. J. Biol. Chem. 289, 21770–21781. doi:10.1074/jbc.M113.541763.

Morris, C. E., Boucher, P.-A., and Joós, B. (2012). Left-Shifted Nav Channels in Injured Bilayer: Primary Targets for Neuroprotective Nav Antagonists? Front. Pharmacol. 3. doi:10.3389/fphar.2012.00019.

Muroi, Y., and Chanda, B. (2009). Local anesthetics disrupt energetic coupling between the voltage-sensing segments of a sodium channel. J. Gen. Physiol. 133, 1–15.

Muyrers, J. P., Zhang, Y., Testa, G., and Stewart, A. F. (1999). Rapid modification of bacterial artificial chromosomes by ET-recombination. Nucleic Acids Res. 27, 1555–1557.

Nagoshi, N., Nakashima, H., and Fehlings, M. G. (2015). Riluzole as a neuroprotective drug for spinal cord injury: from bench to bedside. Mol. Basel Switz. 20, 7775–7789. doi:10.3390/molecules20057775.

Pesti, K., Szabo, A. K., Mike, A., and Vizi, E. S. (2014). Kinetic properties and open probability of α7 nicotinic acetylcholine receptors. Neuropharmacology 81, 101–115. doi:10.1016/j.neuropharm.2014.01.034.

Pittenger, C., Coric, V., Banasr, M., Bloch, M., Krystal, J. H., and Sanacora, G. (2008). Riluzole in the treatment of mood and anxiety disorders. CNS Drugs 22, 761–786.

Ptak, K., Zummo, G. G., Alheid, G. F., Tkatch, T., Surmeier, D. J., and McCrimmon, D. R. (2005). Sodium currents in medullary neurons isolated from the pre-Bötzinger complex region. J. Neurosci. Off J. Soc. Neurosci. 25, 5159–5170. doi:10.1523/JNEUR0SCI.4238-04.2005.

Ragsdale, D. S., McPhee, J. C., Scheuer, T., and Catterall, W. A. (1994). Molecular determinants of state-dependent block of Na+ channels by local anesthetics. Science 265, 1724–1728.

Sheets, M. F., Chen, T., and Hanck, D. A. (2011). Lidocaine partially depolarizes the S4 segment in domain IV of the sodium channel. Pflüg. Arch. -Eur. J. Physiol. 461, 91–97. doi:10.1007/s00424-010-0894-1.

Shen, H., Zhou, Q., Pan, X., Li, Z., Wu, J., and Yan, N. (2017). Structure of a eukaryotic voltage-gated sodium channel at near-atomic resolution. Science 355. doi:10.1126/science.aal4326.

Song, J. H., Huang, C. S., Nagata, K., Yeh, J. Z., and Narahashi, T. (1997). Differential action of riluzole on tetrodotoxin-sensitive and tetrodotoxin-resistant sodium channels. J. Pharmacol. Exp. Ther. 282, 707–714.

Spadoni, F., Hainsworth, A. H., Mercuri, N. B., Caputi, L., Martella, G., Lavaroni, F., et al. (2002). Lamotrigine derivatives and riluzole inhibit INa,P in cortical neurons. Neuroreport 13, 1167–1170.

Starmer, C. F., Grant, A. O., and Strauss, H. C. (1984). Mechanisms of use-dependent block of sodium channels in excitable membranes by local anesthetics. Biophys. J. 46, 15–27. doi:10.1016/S0006-3495(84)83994-6.

Tarnawa, I., Bölcskei, H., and Kocsis, P. (2007). Blockers of voltage-gated sodium channels for the treatment of central nervous system diseases. Recent Patents CNS Drug Discov. 2, 57–78.

Urbani, A., and Belluzzi, O. (2000). Riluzole inhibits the persistent sodium current in mammalian CNS neurons. Eur. J. Neurosci. 12, 3567–3574.

Xie, R.-G., Zheng, D.-W., Xing, J.-L., Zhang, X.-J., Song, Y., Xie, Y.-B., et al. (2011). Blockade of persistent sodium currents contributes to the riluzole-induced inhibition of spontaneous activity and oscillations in injured DRG neurons. PloS One 6, e18681. doi:10.1371/journal.pone.0018681.

Yan, Z., Zhou, Q., Wang, L., Wu, J., Zhao, Y., Huang, G., et al. (2017). Structure of the Nav1.4-β1 Complex from Electric Eel. Cell 170, 470–482.e11. doi:10.1016/j.cell.2017.06.039.

Zamponi, G. W., and French, R. J. (1993). Dissecting lidocaine action: diethylamide and phenol mimic separate modes of lidocaine block of sodium channels from heart and skeletal muscle. Biophys. J. 65, 23–35.

Zarate, C. A., and Manji, H. K. (2008). Riluzole in psychiatry: a systematic review of the literature. Expert Opin. Drug Metab. Toxicol. 4, 1223–1234. doi:10.1517/17425255.4.9.1223.

Zboray, K., Sommeregger, W., Bogner, E., Gili, A., Sterovsky, T., Fauland, K., et al. (2015). Heterologous protein production using euchromatin-containing expression vectors in mammalian cells. Nucleic Acids Res. 43, e102. doi:10.1093/nar/gkv475.

Zhang, H., Zou, B., Du, F., Xu, K., and Li, M. (2015). Reporting sodium channel activity using calcium flux: pharmacological promiscuity of cardiac Nav1.5. Mol. Pharmacol. 87, 207–217. doi:10.1124/mol.114.094789.

Zhang, Y., Buchholz, F., Muyrers, J. P., and Stewart, A. F. (1998). A new logic for DNA engineering using recombination in Escherichia coli. Nat. Genet. 20, 123–128. doi:10.1038/2417.

